# Predicting curvature evolution on biological surfaces from clinical imaging-derived area dilation: a closed-form interpretable framework

**DOI:** 10.64898/2026.05.08.723930

**Authors:** Kameel Khabaz, Charlie Davis, Joseph Pugar, Luka Pocivavsek

## Abstract

Curvature evolution on a deforming surface is governed by the full change in the surface metric, but on biological surfaces captured by serial three-dimensional imaging, only the local area change is observable. The loss of the shear component leaves prediction of curvature evolution underdetermined from imaging alone. On the thoracic aorta, where curvature change marks disease progression, we derive a closed-form equation that predicts the change in integrated Gaussian curvature from the area dilation and initial geometry. The equation combines a conformal term in the area dilation with a leading anisotropy correction from the initial geometry. These two analytic levels, augmented by multi-scale spatial features at neighboring regions and a graph neural network trained on residuals, form a four-level nested predictor. On a synthetic aortic geometry under prescribed isotropic expansion, the equation recovers the analytic coefficient exactly. Across a continuum from pure expansion to pure shear, it holds *R*^2^ ≥ 0.71. On 236 paired thoracic aortic surfaces spanning dissection, aneurysm, traumatic injury, and non-pathologic controls, the equation recovers within-surface curvature change patterns with per-patient median Pearson 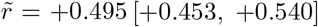 and pooled *R*^2^ = +0.238 [+0.225, +0.250], matching the graph neural network on the same inputs. The residual is a direct measurement of how far the observed growth field departs from conformality.

**Highlights:**

- Closed-form equation predicts aortic curvature change from paired computed tomography scans.
- Recovers analytic predictions exactly on synthetic aortic geometries.
- Anisotropy proxy holds *R*^2^ ≥ 0.71 from pure expansion to pure shear.
- Coefficients tie to geometric mechanisms ensuring interpretability.
- Anisotropy term, computable from one CT, is twice as large on diseased aortas.

## 1 Introduction

The thoracic aorta undergoes complex morphologic changes that mirror disease progression [1–3], yet maximum-diameter criteria for surgical decision-making collapse surface-level information into one scalar and miss the heterogeneous shape changes that drive failed repairs [4]. Aortic surface curvature has been identified as a complementary risk factor to diameter [5–8]. The integrated Gaussian curvature *K* =∫_*R*_ *κ*_*g*_ *dA* over a local region *R* is a natural intrinsic-geometry observable, invariant under isometric reshaping and sensitive only to genuine shape change [9, 10]. We address how *K* changes between two CT scans, given only what clinical imaging makes available.

Patient-specific morphoelastic finite element simulation provides one route to predicting aortic shape evolution [1,11,12], but it requires calibrated material parameters and detailed boundary conditions. Cohort-level descriptors based on integrated Gaussian curvature, including the surface fluctuation *δK* [4, 13] and bending-energy descriptors [14], have been shown to separate aortic outcomes. Neural network models have been applied to related vascular prediction problems, with the trade-off of substantial parameter counts and training-data requirements [15–17]. The present work pursues a complementary direction, deriving a closed-form predictor of the per-patch change in integrated Gaussian curvature directly from the deformation between two CT scans. Such a structure positions the equation as a building block for prospective use as well, including iterated forecasting beyond the second scan and, given a population model of the area dilation field, prediction from a single baseline scan.

The curvature change is set by how the surface metric evolves between the two scans, but imaging recovers only one part of this evolution, the local area change. The shear part is left unobserved, leaving the curvature change underdetermined from imaging alone in the general case. A closed-form bridge exists in a special case, when the deformation is a pure rescaling with no shear; the leading correction outside that case is an anisotropy term computable from the initial geometry alone. Together these give a closed-form predictor of curvature change from imaging-derived inputs [18–22].

We hypothesize that the local change in integrated Gaussian curvature on an aortic surface can be predicted from the area dilation alone, given the initial geometry, when the dilation is small (|*u*| *<* 0.2). On synthetic surfaces under prescribed conformal expansion, where the analytic answer is known, the closed-form fit recovers it exactly. On held-out patient surfaces, the same predictor captures the within-surface pattern of curvature change as well as a flexible graph neural network can on the same inputs. What remains is therefore a measurement of how far observed aortic growth departs from a pure rescaling deformation, not a shortfall of the predictor. The anisotropy term, computable from a single CT scan, is roughly twice as large on diseased as on non-diseased aortic surfaces, providing a geometric marker of shear susceptibility.

## 2 Methods

The methods construct a per-patch predictor of the change in integrated Gaussian curvature Δ*K* between two CT scans. The inputs are the per-patch area dilation *u*, the initial integrated Gaussian curvature *K*_0_, and the initial mean curvature *H*_0_. Section 2.1 describes the patient cohort and the patch-graph construction. Section 2.2 states the closed-form predictor and the origin of each of its terms. Sections 2.3–2.4 organize the features into a nested four-model hierarchy of increasing capacity (Table 1), from the closed-form three-term predictor to a graph neural network used as an upper-capacity reference. Section 2.5 validates each level on idealized synthetic geometries with a known target. Section 2.6 fits the cohort regression on the patient-stratified split, and Section 2.7 reports the regression and tercile classification metrics.

**Table 1:** Feature library and physical meaning, organized by hierarchical model. For each model in the nested hierarchy, the table lists the regression features and a plain-language reading of what each represents. *u* is the per-patch area dilation, *K*_0_ the initial integrated Gaussian curvature, *H*_0_ the initial mean curvature, *κ*_1_, *κ*_2_ the principal curvatures, and Δ, ∇, *W*^*h*^ are operators on the patch graph.

| Term | Physical meaning |
| --- | --- |
| **Liouville model** ($d = 3$) | |
| $\Delta u$ | Patch-graph Laplacian of $u$. The flux of area dilation between a patch and its neighboring patches; the only term derived directly from the conformal Liouville relation. |
| $K_0 u$ | Existing curvature scaled by area dilation. Higher-order term arising from a Taylor expansion of the conformal factor $e^{-2u}$ in the small-$u$ regime. |
| $K_0$ | Initial integrated Gaussian curvature on the patch. Higher-order term included as an initial-geometry baseline, inspired by the lab’s prior $\delta K$ work [4, 13]. |
| **Anisotropy model** ($d = 7$) | |
| $4(H_0^2 - K_0)$ | Squared difference of principal curvatures $(\kappa_1 - \kappa_2)^2$. Shear susceptibility marker that vanishes on umbilic patches and is large on saddle and tubular regions. Combined with $K_0$, recovers the Willmore bending density $\kappa_1^2 + \kappa_2^2$ [14, 28]. |
| $4(H_0^2 - K_0) u$ | Anisotropy times area dilation. Leading non-conformal interaction term motivated by morphoelastic shell theory [20–22]. |
| $\vert \nabla u\vert ^2$ | Dirichlet energy density of $u$. The spatial gradient energy of the area dilation field on the patch graph. |
| $K_0 \vert \nabla u\vert ^2$ | Curvature-coupled gradient energy. |
| **Multiscale model** ($d = 19$) | |
| $B^h u, h \in \{1, 2, 3, 4\}$ | Band-pass area dilation at hop scale $h$, with $B^h u = W^h u - W^{h+1} u$. Captures the spatial structure of $u$ at scale $h$. |
| $B^h K_0, h \in \{1, 2, 3, 4\}$ | Band-pass initial curvature at hop scale $h$. |
| $K_0 (u - W^h u), h \in \{1, 2, 3, 4\}$ | Curvature-coupled multi-hop dilation contrast at scale $h$. |
| **GNN model** |  |
| Multiscale features (input) | Inputs to a graph neural network with a bounded learned residual on top of the Multiscale prediction. Used as an upper-capacity reference for the closed-form fit. |

### 2.1 Patient cohort and patch-graph construction

Two hundred thirty-six patients with paired computed tomography angiography (CTA) scans are identified across four clinical groups: aortic dissection (*Dissection, n* = 103), thoracic aortic aneurysm (*Aneurysm, n* = 55), traumatic aortic injury (*Traumatic, n* = 30), and non-pathologic controls (*Normal, n* = 48). The three pathologic cohorts contribute paired imaging before and after thoracic endovascular aortic repair (TEVAR). The *Normal* cohort contributes paired scans at two timepoints without intervening intervention. The imaging-to-mesh workflow (semi-automated segmentation, surface smoothing, auto-mesh normalization to 0.5 mm element edge, non-rigid registration [23]) follows our prior protocol [1, 4, 13].

Registered initial and final meshes are partitioned into *N* = 800 spatially corresponding patches by centroid-constrained *k*-means with *k*-means++ seeding [24], producing one-to-one correspondence 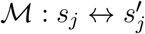 [1]. The patch graph carries edge weights *H*_*ij*_ equal to the boundary length shared between patches *i* and *j* divided by their centroid distance. The discrete Laplacian acts on *u* as Δ*u* = *u* − *Wu*, where *W* = *D*^*−*1^*H* is the row-stochastic random-walk operator and *D* is the row-sum diagonal of *H* [25, 26].

Throughout, lowercase *κ*_*g*_ and *κ*_*h*_ denote pointwise Gaussian and mean curvatures, with *κ*_*g*_ = *κ*_1_*κ*_2_ and *κ*_*h*_ = (*κ*_1_ + *κ*_2_)*/*2, where the principal curvatures *κ*_1_ and *κ*_2_ are estimated on each mesh by Rusinkiewicz’s second-fundamental-form fit [14,27]. Uppercase *K*_0_ and *H*_0_ denote the per-patch initial Gaussian and mean curvatures, computed by area-weighted aggregation of the per-vertex values to the patch level following the convention of our prior work [1, 4]. The target Δ*K* = *K*(*t*_1_) − *K*(*t*_0_) is the change in *K* between scans, and each patch also carries the log area ratio 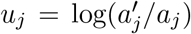. Training requires *κ*_*g*_ on both scans. Inference on a new patient needs only *u* from registration and the initial-scan curvatures {*K*_0_, *H*_0_, *κ*_1_, *κ*_2_}, since per-patient demeaning sets∑ _*j*_ Δ*K*_*j*_ = 0 on each surface. The per-surface root-mean-square |*u*| is 0.10 ± 0.04, within the small-*u* regime of the linearized Liouville expansion.

### 2.2 Closed-form predictor

Three imaging-derived measurements are available per patch: the area dilation *u* = log(*a*^*′*^*/a*), the initial integrated Gaussian curvature *K*_0_, and the initial mean curvature *H*_0_. Features are constrained to be functions of these three quantities. The closed-form predictor reads

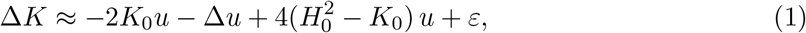

where Δ*u* is the patch-graph Laplacian of *u* and *ε* is the residual that absorbs the unobserved non-conformal shear. Each term has a distinct origin. The patch-graph Laplacian Δ*u* is derived directly from the conformal Liouville relation between two metrics related by 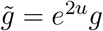 [18, 19,25,26]; under purely conformal evolution and patch integration, it is the exact source for the change in integrated Gaussian curvature on the patch. The *−2K*_0_*u* term is a higher-order correction obtained by Taylor-expanding the conformal factor *e*^*−*2*u*^ in the small-*u* regime (|*u*| *<* 0.2 on the cohort). The bare *K*_0_, included as an additional regression feature in the model, is a higher-order initial-geometry baseline inspired by the lab’s prior *δK* work [4, 13]. The 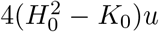 term is the leading non-conformal correction motivated by morphoelastic shell theory [20–22]; the anisotropy factor 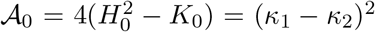 vanishes on umbilic patches and is large on saddle and tubular regions, serving as a shear susceptibility marker. Combined with *K*_0_, it recovers the Willmore bending density 𝒜_0_ + 2*K*_0_ = *κ*^2^ + *κ*^2^ [14, 28].

### 2.3 Per-patch geometric feature library

The regression features are organized as a nested four-model hierarchy of increasing capacity (Table 1). The **Liouville** model (*d* = 3) takes the three terms appearing in Eq. (1), with the bare *K*_0_ added as an initial-geometry baseline. The **Anisotropy** model (*d* = 7) augments with the shear susceptibility proxy 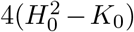, its coupling 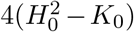· *u*, the Dirichlet gradient-energy density |∇*u*|^2^, and a curvature-coupled form *K*_0_ · |∇*u*|^2^, each evaluated on the patch graph [25]. The **Multiscale** model (*d* = 19) adds twelve multi-hop band-pass features that capture the spatially non-local response of Δ*K* to *u*. The band-pass operator *B*^*h*^*x* = *W*^*h*^*x* − *W*^*h*+1^*x* isolates the spatial structure of *x* at hop scale *h*, where *W* is the row-normalized graph-Laplacian smoother on the patch graph. The twelve features comprise band-pass *u* at four hop scales *h* ∈ {1, 2, 3, 4}, band-pass *K*_0_ at the same four scales, and the curvature-coupled multi-hop dilation contrast *K*_0_ · (*u* − *W*^*h*^*u*) at the same four scales [25]. The **GNN** model adds a graph-aware non-linear correction on top of Multiscale, with architecture and training described in Section 2.4. Term-level physical readings are given in Table 1; the seven sparse-selected coefficients of the headline equation are reported in Table 3.

### 2.4 Nested family of predictive models

The Liouville, Anisotropy, and Multiscale models are fit by ridge regression. The ridge penalty *α* is chosen on held-out validation patients (split defined in Section 2.6), and 95% confidence intervals come from *B* = 1000 patient-stratified bootstrap resamples [29]. The headline sparse equation (Table 3) reports the seven Multiscale coefficients with the largest magnitude after *𝓁*_2_ regularization at *N* = 800, with bootstrap 95% CIs reported alongside each.

The GNN model is a three-layer message-passing graph neural network with 128 hidden units per layer [30–32], trained on the residual 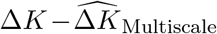with roughly 2.2× 10^5^ tunable parameters. The output correction is bounded at 4*σ* of the Multiscale training residual via tanh scaling, so the GNN cannot deviate arbitrarily from the closed-form prediction. It serves as a flexible upper-envelope reference rather than as the recommended interpretable predictor.

All models share a per-patient pipeline. The dilation *u* is z-scored within each patient, and Δ*K* is demeaned within each surface to neutralize per-surface noise heterogeneity without leakage. A sensitivity sweep across three normalization regimes (per-patient, per-cohort, global) yields pooled test *R*^2^ within 0.016 of one another (Suppl. Table ST3). All coefficients are reported on the demeaned, z-scored scale.

### 2.5 Validation on synthetic conformal growth

The equation form is validated on a synthetic aortic geometry, a cylindrical descending segment joined to a half-toroidal arch with 600 patches. Four prescribed deformation modes are applied at five extents *ε* ∈ {0.05, 0.10, 0.20, 0.40, 0.60}.

Mode A is a uniform conformal radial expansion, the strict-conformal sanity case. Mode B is a Gaussian radial bulge at the arch apex (*σ* = 15 mm) where *K*_0_ *>* 0, designed to test the Liouville source *K*_0_ ·*u*. Mode C is a narrow non-axisymmetric bulge in the cylindrical segment (*σ* = 8 mm) where *K*_0_ ≈ 0, designed to silence *K*_0_·*u* and make the second-order Liouville term *u*·Δ*u* the leading contribution. Mode D consists of four amplitude-weighted non-axisymmetric bulges along the cylindrical segment at sigmas {4, 8, 16, 28} mm, chosen to span the four band-pass hop scales of the patch graph.

Each mode is fit separately with the same Liouville/Anisotropy/Multiscale ridge pipeline. The synthetic surfaces do not inherit per-patient z-scoring or per-surface demeaning of Δ*K*. Those transforms are reserved for the cohort fits.

### 2.6 Cohort-stratified data partition

All quantitative claims rest on a single 70%/15%/15% patient-stratified split of the 236-patient cohort, with 164 training patients, 35 validation patients, and 37 test patients. The split preserves clinical-group proportions (Dissection, Aneurysm, Normal, Traumatic) and is fixed across all models, resolutions, and bootstrap resamples. Leave-one-out and repeated cross-validation are avoided in the main pipeline because their dependence structure complicates bootstrap CIs. A leave-one-cohort-out sensitivity analysis is reported in Suppl. Section 2.

Bootstrap 95% percentile CIs on the pooled test *R*^2^ come from *B* = 1000 patch-level resamples of the held-out test patches with replacement [29]. Bootstrap CIs on the ridge coefficients *β* come from *B* = 1000 patch-level resamples of the training set, refitting the ridge on each resample. The per-patient Pearson *r* is summarized as the across-patient median with the 25th–75th percentile interquartile range. Ridge *α* is selected on validation patients by grid search over *α* ∈ {10^*−*3^, 10^*−*2^, …, 10^5^} and fixed before test evaluation.

### 2.7 Regression and tercile-classification metrics

The primary regression metric is the pooled test *R*^2^, computed across all 37 × 800 = 29,600 test patches on per-patient demeaned Δ*K*. The pooled metric is complemented by the per-patient Pearson correlation *r*, reported as median and IQR over 37 test patients. The per-patient framing quantifies the clinically relevant within-surface pattern match [15] and is robust to the order-of-magnitude variation in Δ*K* noise scale across patients.

The primary classification metric is per-patient tercile agreement. Each test patch is labeled bottom, middle, or top tercile of the truth’s Δ*K* within its own surface, and the prediction is binned by the same cut-points. The resulting three-class problem is balanced, with chance accuracy 1*/*3 and chance *κ* = 0. Macro *F*_1_, accuracy, Cohen’s *κ*, and the 3 × 3 confusion matrix are reported. Tercile framing is preferred over |*z*|-thresholded and quartile-extreme alternatives,which inflate *κ* through middle-class dominance. A residual Moran’s *I* is also reported [33, 34], with local indicators of spatial association [35] reported in Suppl. Section 12. Models are also compared against five non-physical baselines defined in Suppl. Section 7: a no-growth predictor, a per-patient mean predictor, a linear-history *k*-NN extrapolation [15], and tuned random-forest and XGBoost regressors on the Multiscale feature matrix.

## 3 Results

### 3.1 Synthetic validation across deformation modes

On the synthetic aortic geometry, four prescribed deformation modes test the contribution of each successive model layer (Fig. 1). Mode A, a uniform conformal expansion, reaches *R*^2^ = +1.000 on all three models at every extent *ε* ∈ {0.05, 0.10, 0.20, 0.40, 0.60} and saturates the strict-conformal sanity floor.

**Figure 1:**
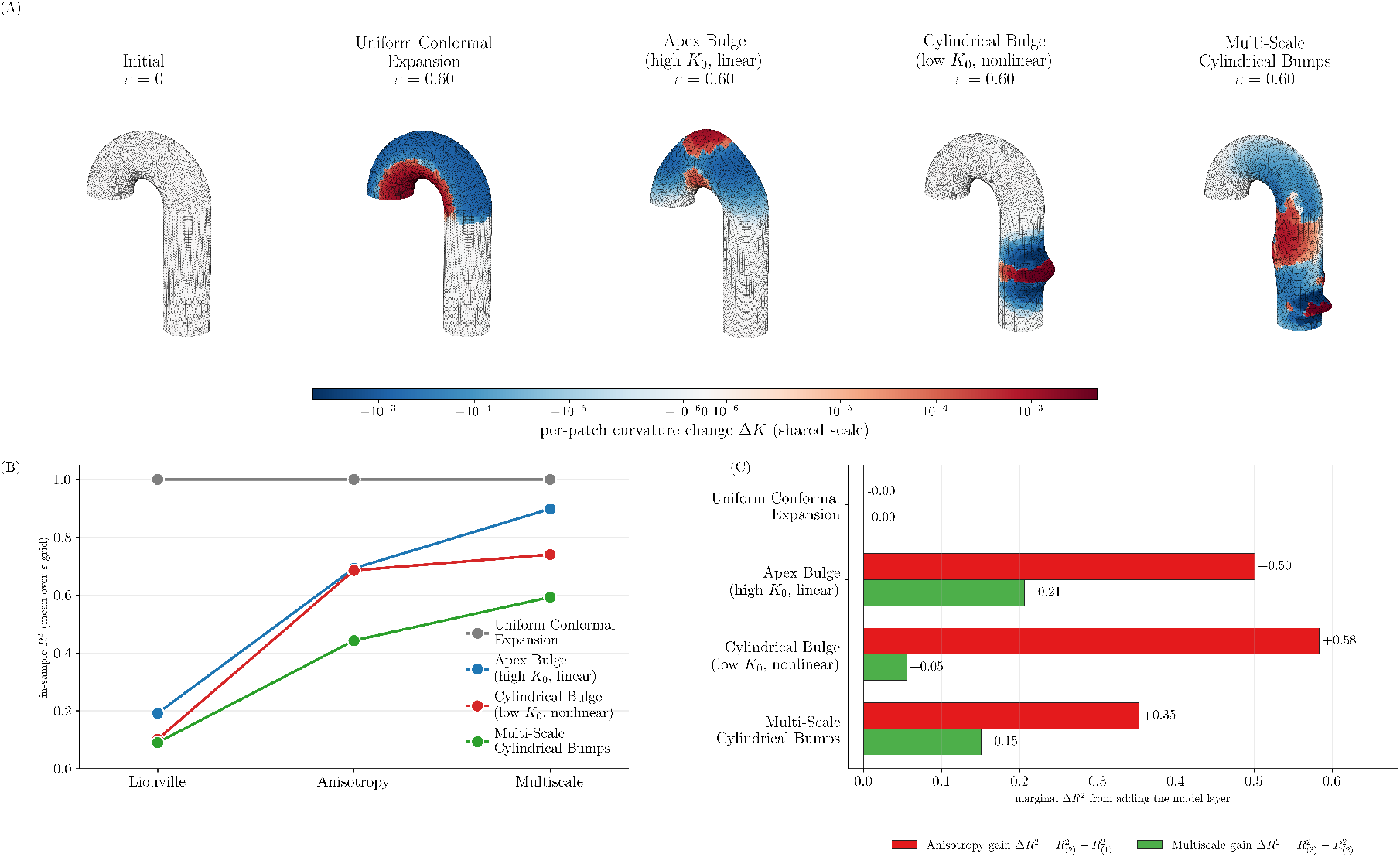
Validation of Δ*K* prediction on synthetic aortas across four deformation modes. The Liouville, Anisotropy, and Multiscale models are fit on a synthetic aortic geometry under four prescribed deformation modes at deformation magnitude *ε* = 0.60. **(A)** Initial geometry and the four deformed surfaces, colored on a shared symlog Δ*K* scale. The deformation modes are uniform conformal expansion, a Gaussian radial bulge at the apex, where *K*_0_ *>* 0 and the linearised Liouville source *K*_0_ ·*u* is the dominant contribution to Δ*K*; a narrow non-axisymmetric radial bulge in the cylindrical descending segment, placed where *K*_0_ ≈0 to silence *K*_0_· *u* so that the second-order term *u* ·Δ*u* becomes the leading non-trivial contribution; and four non-axisymmetric bulges along the same cylindrical segment at sigmas {4, 8, 16, 28} mm spanning the four band-pass hop scales of the patch graph. **(B)** In-sample *R*^2^ per mode across the model hierarchy (Liouville: three features; Anisotropy: seven; Multiscale: nineteen). Each mode follows a distinct path. Mode A saturates at unity. Mode B gives *R*^2^ = +0.19→ +0.69→ +0.90. Mode C drops from *R*^2^ = +0.16 at *ε* = 0.05 to +0.06 at *ε* = 0.60 on Liouville and climbs from +0.65 at *ε* = 0.05 to +0.72 at *ε* = 0.60 on Anisotropy, where the quadratic feature |∇*u*|^2^ captures the nonlinear *u*·Δ*u* content that the linear triple drops. Mode D gives *R*^2^ = +0.09 → +0.44 → +0.59, with the largest gain entering at the Multiscale layer through the multi-hop band-pass features 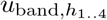 . **(c)** Marginal Δ*R*^2^ added by each layer per mode, exposing which model addition earns each gain: Anisotropy adds +0.46 on B and +0.69 on C; Multiscale adds +0.10 on D where Anisotropy added only +0.29. Modes B, C, and D therefore independently motivate the three non-trivial model layers in the hierarchy.

Modes B, C, and D are non-conformal cases, each motivating one of the three non-trivial layers in the closed-form hierarchy. The focal apex bulge of Mode B (where *K*_0_ *>* 0) gives mean *R*^2^ values across the *ε* grid of +0.19 (Liouville), +0.69 (Anisotropy), and +0.90 (Multiscale). The anisotropy proxy 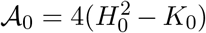 recovers the non-conformal residual that the linearized triple misses. The narrow non-axisymmetric bulge of Mode C in the cylindrical region (where *K*_0_ ≈ 0) silences the linear *K*_0_ ·*u* source. Liouville *R*^2^ drops from +0.16 at *ε* = 0.05 to +0.06 at *ε* = 0.60, while Anisotropy climbs from +0.65 at *ε* = 0.05 to +0.72 at *ε* = 0.60 because the quadratic Dirichlet term |∇*u*|^2^ in the second-order Liouville expansion remains effective when the linear source vanishes. The four multi-scale non-axisymmetric bulges of Mode D give mean *R*^2^ values of +0.09, +0.44, and +0.59 across Liouville, Anisotropy, and Multiscale. The largest gain enters at the Multiscale layer through the multi-hop band-pass features 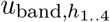, which span the four bump scales that no single-hop feature resolves.

The nesting Liouville ≤ Anisotropy ≤ Multiscale is *ε*-invariant on every mode. The equation form is therefore anchored on analytically distinguishable cases before the same fitting pipeline is applied to clinical data.

A conformal–non-conformal mixing sweep on the same synthetic aorta tests the robustness of the anisotropy correction to deviations from the conformal special case (Fig. 2). The deformation is the convex combination *α u*_conf_ +(1−*α*) *u*_aniso_ of a global conformal expansion (*ε*_*A*_ = 0.10) and a focal anisotropic stretch at the apex (*ε*_*B*_ = 0.10), at *α* ∈ {0, 0.25, 0.5, 0.75, 1.0}. The Liouville *R* increases from +0.281 at pure anisotropic stretch (*α* = 0) to +1.000 at pure conformal expansion (*α* = 1), tracing the failure of the strict conformal predictor as the deformation acquires deviatoric content. The Anisotropy model holds *R*^2^ ≥ +0.747 across the entire mixing spectrum, with the shear gap 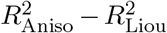collapsing super-linearly in *α* (maximum deviation of the observed shear gap from the linear-interpolation baseline reaches 207% of the endpoint difference at *α* = 0.25, with the gap closed to ≤ +0.001 at *α* ≥ 0.5). The anisotropy proxy 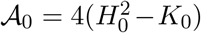 therefore acts as a mode-robust lower bound on Δ*K* across the full conformal– non-conformal spectrum, not only at the pure-mode endpoints.

**Figure 2:**
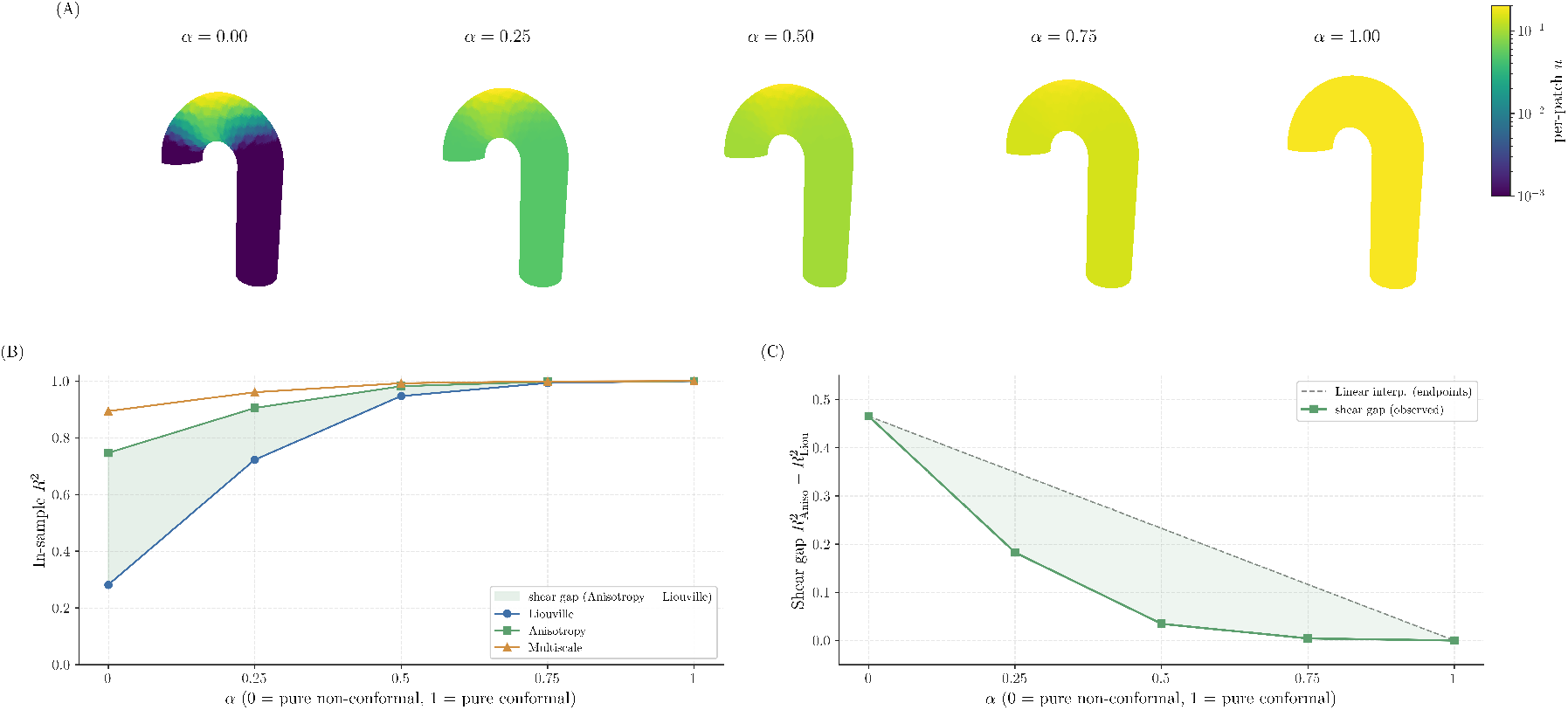
Conformal–non-conformal superposition test on the idealized aorta. On the same synthetic geometry as Fig. 1, the deformation is the convex combination *α* · *u*_conf_ + (1 − *α*) · *u*_aniso_ of a global conformal expansion (*ε*_*A*_ = 0.10) and a focal anisotropic stretch at the arch apex (*ε*_*B*_ = 0.10), at *α* ∈{0, 0.25, 0.5, 0.75, 1.0} . (Left) In-sample *R*^2^ of the Liouville (blue), Anisotropy (red), and Multiscale (green) closed-form models versus mixing fraction. The Liouville *R*^2^ rises from +0.281 at pure non-conformal stretch to +1.000 at pure conformal expansion; the Anisotropy *R*^2^ stays at or above +0.747 across the entire sweep. The red shaded band is the shear gap 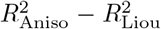. (Right) The shear gap collapses super-linearly in *α* (red, observed) compared to the linear-interpolation baseline (dashed grey), with maximum deviation 215% of the endpoint difference at *α* = 0.25. The super-linearity reflects the spatial dominance of global conformal dilation over focal anisotropic stretch in pooled variance: even a partial conformal share suffices to close most of the shear gap. The anisotropy term 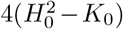 therefore acts as a mode-robust lower bound on Δ*K* across the full conformal–non-conformal spectrum, not only at the pure-mode endpoints.

### 3.2 Cohort prediction across the model hierarchy

On the cohort, per-patient median Pearson 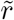 increases from +0.335 (Liouville) to +0.495 (Multiscale) across the closed-form hierarchy on the 37 held-out test surfaces, a 48% improvement in within-surface pattern match (Fig. 3, Table 2). Pooled test *R*^2^ increases over the same range from +0.117 to +0.238.

**Table 2:** Liouville, Anisotropy, Multiscale, and GNN model performance on the held-out test cohort. Pooled train, validation, and test *R*^2^ together with the patient-stratified bootstrap 95% confidence interval, the per-patient median Pearson 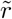 with interquartile range, and the per-patient tercile Cohen’s *κ*. Model dimensionality *d* counts the number of regression features. Bootstrap CIs are computed from *B* = 1000 resamples of the 37 held-out test patients.

| Model | $d$ | train $R^2$ | val $R^2$ | test $R^2$ [95% CI] | per-patient $r$ med [IQR] | tercile $\kappa$ |
| --- | --- | --- | --- | --- | --- | --- |
| Liouville | 3 | +0.120 | +0.136 | +0.117 [+0.107, +0.127] | +0.335 [+0.272, +0.366] | +0.138 |
| Anisotropy | 7 | +0.140 | +0.156 | +0.135 [+0.124, +0.145] | +0.358 [+0.301, +0.418] | +0.154 |
| Multiscale | 19 | +0.256 | +0.273 | +0.238 [+0.225, +0.250] | +0.495 [+0.453, +0.540] | +0.241 |

**Table 3:** Sparse top-7 coefficients of the Multiscale model fit. Each row reports a feature name, its physical meaning, the bootstrap mean *β*, the bootstrap 95% confidence interval, and the sign. The linearized Liouville source *K*_0_· *u*, the initial surface anisotropy 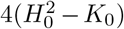, the coupling of anisotropy to strain, 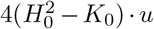, and the initial Gaussian curvature *K*_0_ carry the largest magnitudes after rescaling and bootstrap-stable positive signs. These four coefficients are the bootstrap-significant survivors of the sparse selection step described in Section 2.4; the three remaining sparse-selected terms have confidence intervals that bracket zero on the 800-patch fit and are reported in the table for completeness.

| Term | Physical meaning | $\beta$ | 95% CI | sign |
| --- | --- | --- | --- | --- |
| $4(H_0^2 - K_0)$ | Initial-surface anisotropy (shear proxy) | +9.75e-02 | [+9.11e-02, +1.04e-01] | + |
| $K_0$ | Initial Gaussian curvature | +8.75e-02 | [+8.12e-02, +9.38e-02] | + |
| $K_0 \cdot u$ | Linearized Liouville source | +7.07e-02 | [+6.39e-02, +7.79e-02] | + |
| $4(H_0^2 - K_0) \cdot u$ | Anisotropy–strain coupling | +6.57e-02 | [+5.87e-02, +7.29e-02] | + |
| $\Delta u$ | Discrete Laplacian of area dilation | -3.62e-04 | [-5.62e-04, -1.51e-04] | – |
| $\vert \nabla u\vert ^2$ | Dirichlet energy density of area dilation | -1.32e-04 | [-3.31e-04, +6.19e-05] | – |
| $K_0 \vert \nabla u\vert ^2$ | Curvature-coupled Dirichlet energy | +3.37e-05 | [-3.49e-04, +4.18e-04] | + |
| intercept |  | -5.69e-19 |  |  |

**Figure 3:**
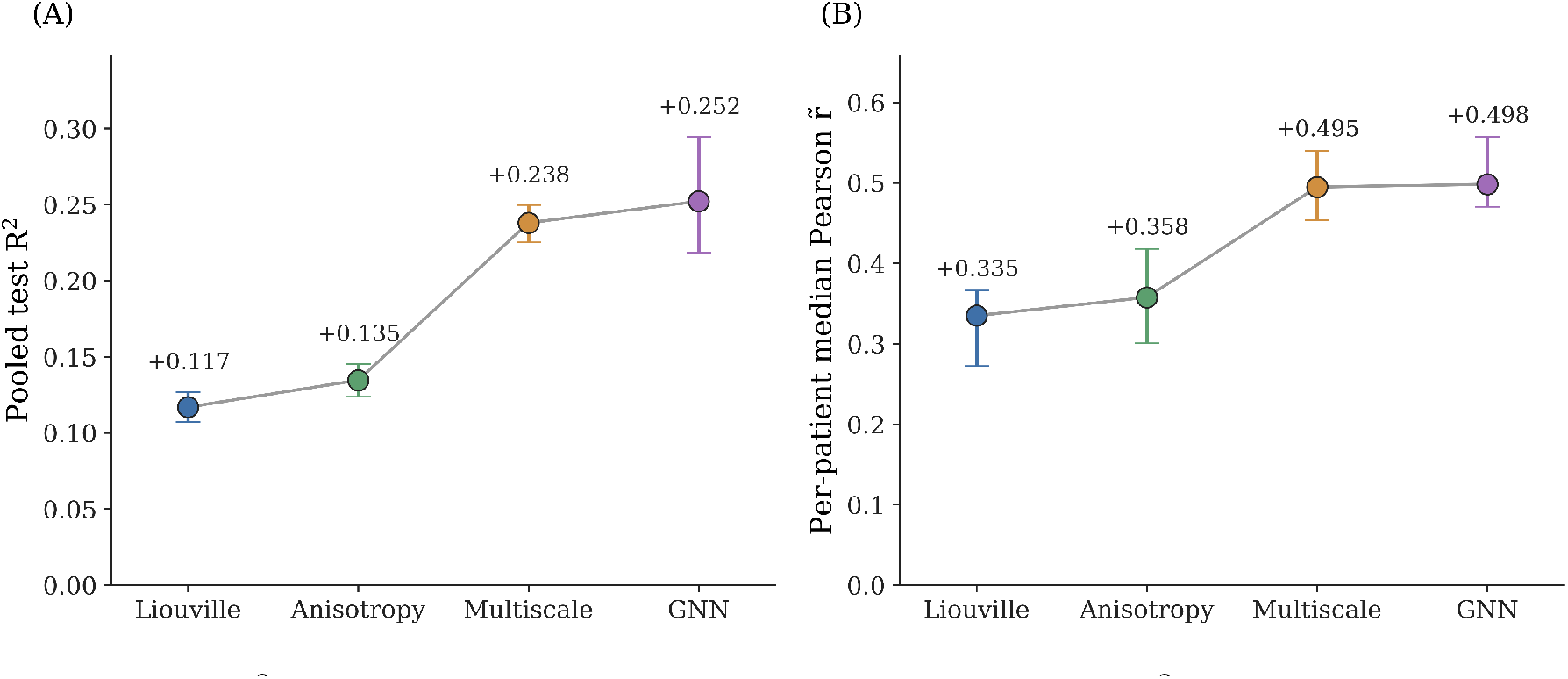
Test *R*^2^ across the nested model hierarchy. Pooled test *R*^2^ on the 37 held-out patients increases monotonically from +0.117 for the Liouville model (linearized Liouville source, *d* = 3) to +0.135 for Anisotropy (adding the non-conformal 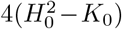 correction, *d* = 7) to +0.238 for Multiscale (adding multi-hop band-pass features, *d* = 19) and to +0.251 for the GNN ceiling (bounded-residual GNN, *d* = 19). Dots are bootstrap mean test *R*^2^ at each level; error bars show the patient-stratified bootstrap 95% confidence intervals over *B* = 1000 resamples; the right panel shows the per-patient median Pearson 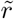 with interquartile range. The Liouville-to-Anisotropy gain is small (Δ*R*^2^ = +0.018); the Anisotropy-to-Multiscale gain (Δ*R*^2^ = +0.103) is the dominant out-of-sample improvement; GNN sits at the Multiscale ceiling within bootstrap CIs (Δ*R*^2^ = +0.013).

The Liouville model reaches 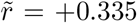, *R*^2^ = +0.117 [+0.107, +0.127]. The Anisotropy model reaches 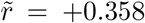, *R*^2^ = +0.135 [+0.124, +0.145]. The Multiscale model reaches 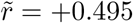, *R*^2^ = +0.238 [+0.225, +0.250], *κ* = +0.241. The GNN model reaches 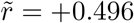,*R*^2^ = +0.251 [+0.236, +0.265], *κ* = +0.261.

The gain from Liouville to Anisotropy is Δ*R*^2^ = +0.018, small but non-zero on the cohort. The case for the anisotropy term rests on sign and significance evidence (Section 3.3) together with this modest *R*^2^ gain. The gain from Anisotropy to Multiscale is Δ*R*^2^ = +0.103, the dominant out-of-sample improvement, reflecting spatial non-locality of Δ*K*’s response to *u* that the local Liouville triple cannot represent without multi-hop band-pass features.

The gap from Liouville to Anisotropy traces to non-conformal modes, not to a nonlinearly conformal residual. Augmenting the Anisotropy model with the higher-order conformal *K*_0_*u*^2^ and *u* Δ*u* terms adds no further *R*^2^ on the cohort fit, which pins the gap on shear that the anisotropy proxy 𝒜_0_ is designed to capture.

The GNN model (test *R*^2^ = +0.251 [+0.236, +0.265]) sits at the Multiscale ceiling within bootstrap CIs (Δ*R*^2^ = +0.013), so the bounded graph-aware correction adds little out-of-sample headroom over the closed-form prediction. The headline is therefore the closed-form Multiscale equation of Section 3.3, and GNN stands as a non-linear ceiling estimate rather than a recommended predictor.

The headline coefficient signs and the cross-level ordering Liouville *<* Anisotropy ≲ Multiscale hold across patch-graph resolutions *N* ∈ {100, 200, 400, 800, 1000} within ±0.04 at every level (Suppl. Section 1). They also hold under a leave-one-clinical-group-out sensitivity in which every held-out group, including the smallest two (Normal, *n* = 48, and Traumatic, *n* = 30), retains positive Multiscale test *R*^2^ (Suppl. Section 2). The Multiscale test *R*^2^ = +0.238 exceeds every non-physical baseline the two-timepoint cohort admits, including the linear-history extrapolation of [15] (*R*^2^ = +0.059) and tuned random-forest and XGBoost regressors on the same features (*R*^2^ = +0.221 and +0.212, with bootstrap CIs non-overlapping the Multiscale CI). The full baseline panel is in Suppl. Section 7.

### 3.3 Coefficients of the linearized Liouville predictor

The four leading coefficients of the headline equation sit at bootstrap-mean *β* ∈ [+0.066, +0.098] with 95% CIs strictly above zero, two orders of magnitude larger than the gradient energy and Laplacian terms (Fig. 4, Table 3). The prediction is organized around the linearized Liouville source *K*_0_ · *u* and the initial surface anisotropy 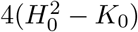,

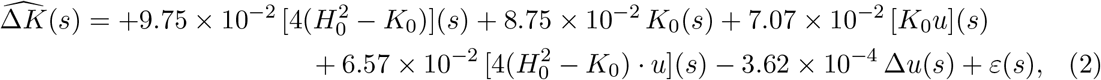

where each coefficient is the bootstrap mean (Table 3).

**Figure 4:**
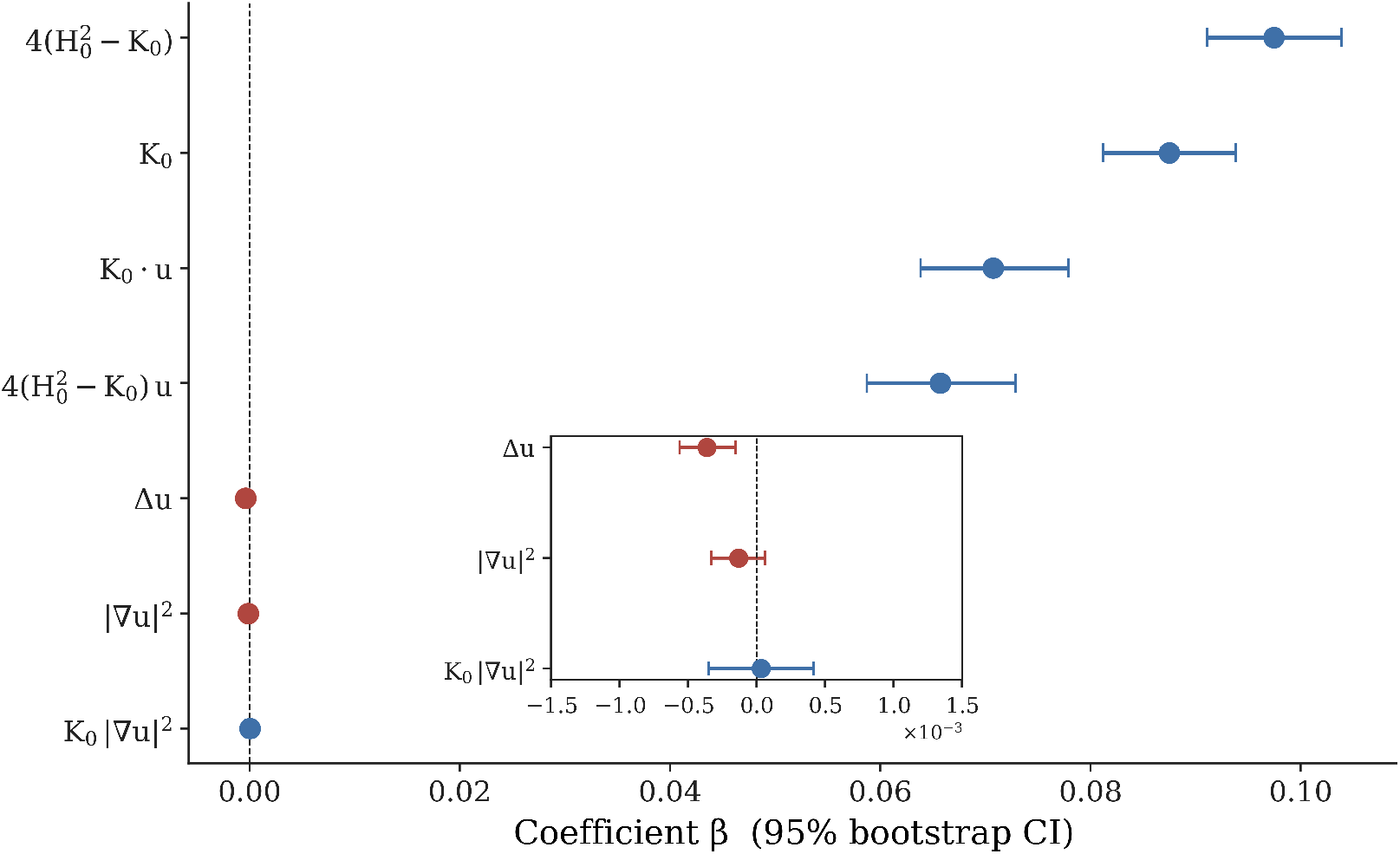
Sparse top-7 coefficient forest plot for the Multiscale model on the held-out test cohort. (*n* = 37 patients, *N*_patches_ = 29,600). The seven coefficients of the headline equation reported in Table 3 are shown as bootstrap mean *β* with 95% confidence intervals from *B* = 1000 patient-stratified resamples. Blue dots mark positive coefficients, red dots negative; coefficients whose 95% interval excludes zero are bootstrap-significant and reported in Eq. (2). The inset zooms into *β* ∈ [−0.005, +0.005] to resolve the three sub-leading terms (Δ*u*, |∇*u*|^2^, *K*_0_ |∇*u*|^2^) whose magnitudes are two orders of magnitude smaller than the local physics quartet on the main axis. The linearized Liouville source *K*_0_ ·*u* and the initial surface anisotropy _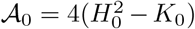_enter with the largest magnitudes after rescaling, with positive signs stable under bootstrap, providing the empirical evidence for both the conformal Liouville term and the non-conformal correction from shell theory.

The positive fitted 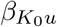 is opposite the strict-conformal −2*K*_0_*u* in Eq. (1). On the idealized arch the same fit recovers the analytic Liouville coefficient under uniform conformal expansion and flips positive under a focal non-conformal bulge (Suppl. §4), placing the cohort in the non-conformal regime. Both anisotropy terms enter at magnitudes comparable to *K*_0_ · *u* with bootstrap intervals excluding zero, confirming that the non-conformal shell correction [20–22,36] is a necessary contributor. The Laplacian Δ*u* enters with a small negative coefficient whose 95% bootstrap CI excludes zero on the 800-patch fit, but with magnitude two orders below the local physics quartet. The two remaining sparse-selected coefficients (|∇*u*|^2^ and *K*_0_ · |∇*u*|^2^) fail bootstrap significance and are dropped from the displayed equation. Full coefficients are listed in Table 3.

The prediction is dominated by pointwise products *K*_0_ · *u* and 𝒜_0_ · *u* together with the initial geometry terms *K*_0_ and 𝒜_0_, rather than by spatial derivative information. This dominance is consistent with Eq. (1). The Laplacian Δ*u* has unit coefficient in raw units, and the per-surface z-scoring of *u* (Var(*u*) ≫ Var(Δ*u*)) suppresses its scaled coefficient.

An *𝓁*_1_-sparsity sweep over the Multiscale feature matrix makes the equation-collapse claim explicit (Suppl. Section 6). The seven-coefficient equation Eq. (2) sits at *R*^2^ = +0.135. The +0.103 gap to the Multiscale headline +0.238 is contributed jointly by the twelve multi-hop band-pass features, none of which individually survives L1 sparsification. Among the seven sparse-selected coefficients, the four leading local physics terms (*K*_0_, *K*_0_*u*, A_0_, A_0_*u*) carry the entire sparse-equation *R*^2^ to within 5 × 10^*−*4^ [37]. The four-term *minimal headline equation* therefore reads

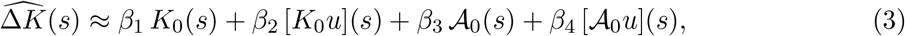

with the four bootstrap-significant coefficients of Table 3. The predictive content lives entirely in the pointwise products of *K*_0_ and 𝒜_0_ with *u*. Pooling all 29,600 held-out patches after per-surface SD normalization collapses the four clinical-group clouds onto a single master curve (Fig. 5) with pooled *r* = +0.480 [+0.467, +0.493] and identity-slope *α* = 0.259 [+0.250, +0.268]. The under-unity slope is the visual signature of the residual variance that no function of *u* alone can capture, the gap between the kinematic observable and the curvature change left by the unobserved deviatoric shear. We refer to this as the *shear gap* throughout, and quantify it explicitly in Section 4 via the 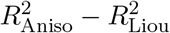 envelope on synthetic deformations.

**Figure 5:**
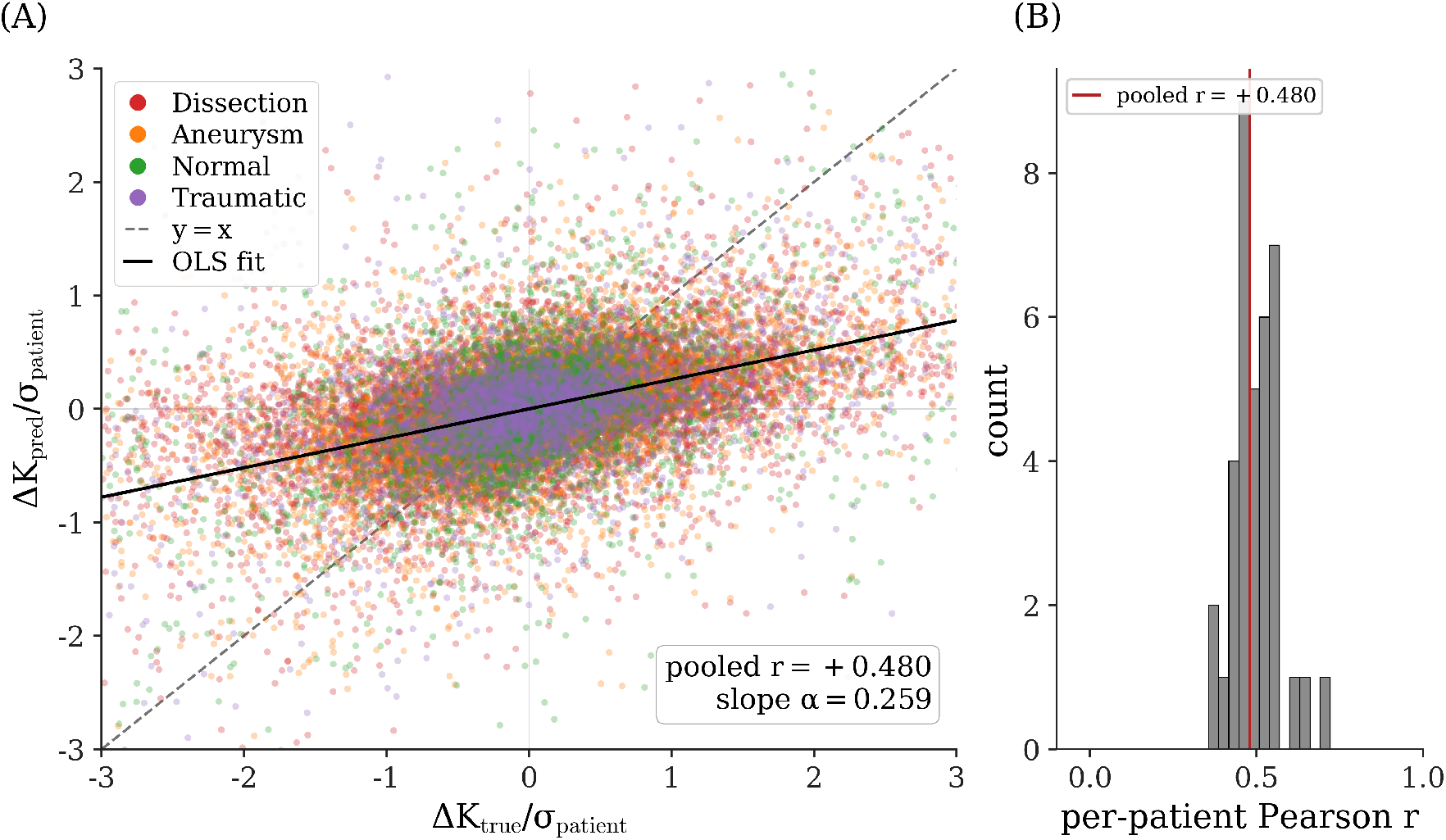
Master curve of predicted versus true curvature change across the held-out test cohort. Each point is a single patch on one of the 37 held-out aortic surfaces (*N*_patches_ = 29,600 total); both axes are normalized by the per-patient standard deviation *σ*_patient_ of the true curvature change Δ*K* so that surfaces of widely differing magnitudes collapse onto a common scale. The interpretable Multiscale ridge equation, fit only on training-cohort patches, is evaluated on every held-out surface and the resulting (Δ*K*_true_, Δ*K*_pred_) pairs are pooled. The pooled Pearson correlation is *r* = +0.480 (95% bootstrap CI: [+0.467, +0.493]); the identity-test ordinary-least-squares slope is *α* = 0.259 (95% CI: [+0.250, +0.268]). Points are colored by clinical group (Dissection, Aneurysm, Normal, Traumatic); the dashed grey line is the identity *y* = *x* and the solid black line is the OLS best fit. The right panel shows the distribution of per-patient Pearson *r* across the 37 held-out surfaces (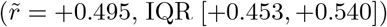, IQR [+0.453, +0.540]), with the pooled value marked in red. The collapse of four clinical-group clouds onto one shared trend demonstrates that the headline interpretable equation captures a cross-patient consistent mapping from area dilation to curvature change rather than a per-patient idiosyncrasy.

### 3.4 Cohort-conditional coefficient magnitude

Refitting the headline equation separately within each clinical group on the train+val patient subset preserves the equation form while separating its magnitude across diseased and non-diseased geometries (Fig. 9). The four leading local physics coefficients (𝒜_0_, *K*_0_, *K*_0_*u*, 𝒜_0_*u*) are uniformly positive across all four groups with bootstrap 95% CIs strictly above zero, confirming that the predictor functions on physiologic, dissection, aneurysmal, and traumatic geometries from a single equation form. The two anisotropy-driven coefficients separate diseased from non-diseased cohorts. The Aneurysm cohort gives *β*_*𝒜*0_= +0.163 [+0.145, +0.179] and Dissection gives +0.140 [+0.127, +0.152], against +0.097 [+0.080, +0.109] on Traumatic and +0.076 [+0.066, +0.086] on Normal, roughly half the diseased magnitude. The separation is sharper on the coupled term 𝒜_0_ ·*u*, where the Aneurysm *β* = +0.127 is ∼ 3.6× the Normal *β* = +0.035. The cross-cohort ordering Aneurysm ≳ Dissection ≫ Traumatic ≳ Normal is preserved across all four leading panels, identifying the non-conformal correction as quantitatively larger in pathologic remodeling than in physiologic or post-traumatic geometries.

### 3.5 Within-surface prediction quality

The per-patient median Pearson 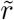 is +0.495 (IQR [+0.453, +0.540]) across the 37 held-out patients (Fig. 6). The narrow IQR indicates that prediction quality is consistent across patients.

**Figure 6:**
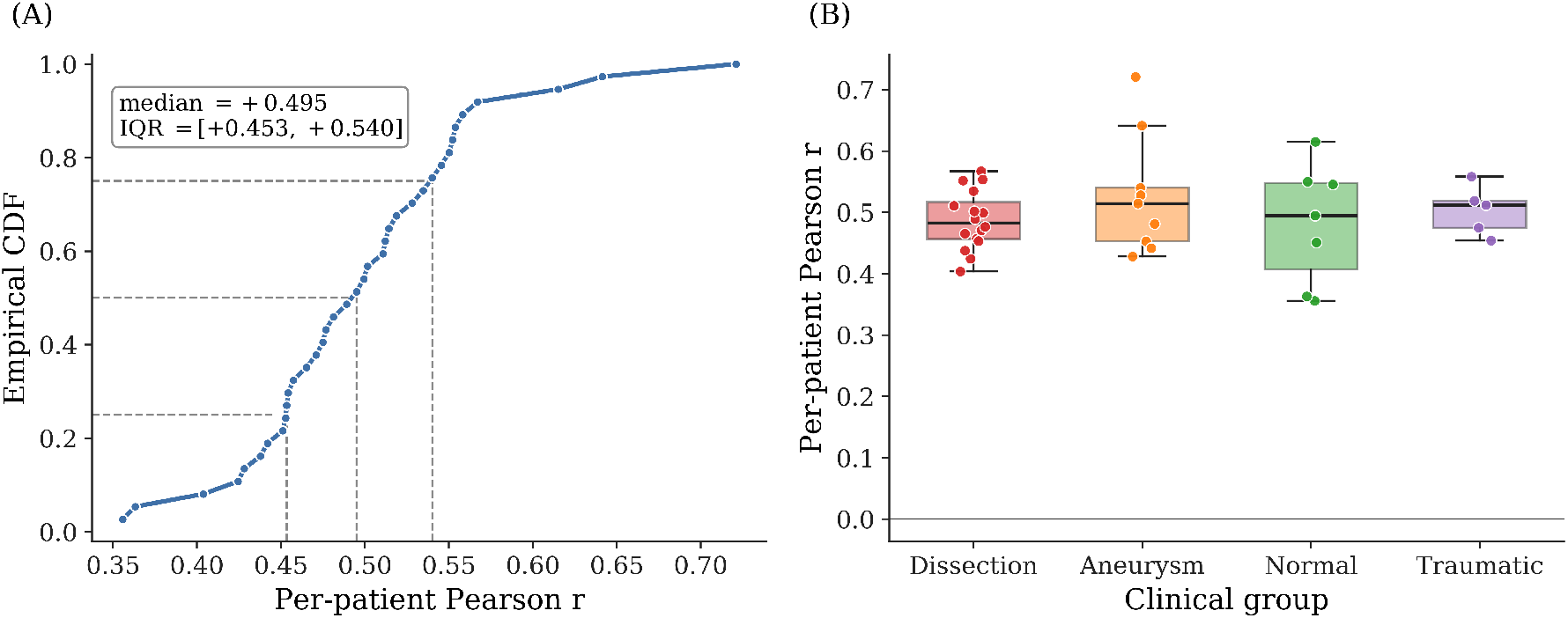
Per-patient Pearson *r* across the held-out cohort. The empirical cumulative distri-bution of per-patient Pearson *r* (left panel) and the cohort boxplot (right panel) summarize prediction quality on each of the 37 held-out test patients, with 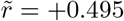 and interquartile range [+0.453, +0.540]. Per-patient heterogeneity reflects per-surface noise scale variation rather than systematic model failure on any clinical cohort, motivating the per-patient framing of the regression metric.

The best test surface (Aneurysm) reaches *r* = +0.721 and per-patient *R*^2^ = +0.373. The median surface (Dissection) reaches *r* = +0.499, *R*^2^ = +0.249. The worst surface (Normal) reaches *r* = +0.356, *R*^2^ = +0.096 (Fig. 7). The best and median surfaces visibly recover sign and spatial localization of large ±Δ*K* patches. The worst preserves localization but compresses magnitude toward zero. The full spatial structure of *u*, true Δ*K*, predicted 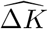, and residual fields on the same three surfaces is shown in Fig. 8.

**Figure 7:**
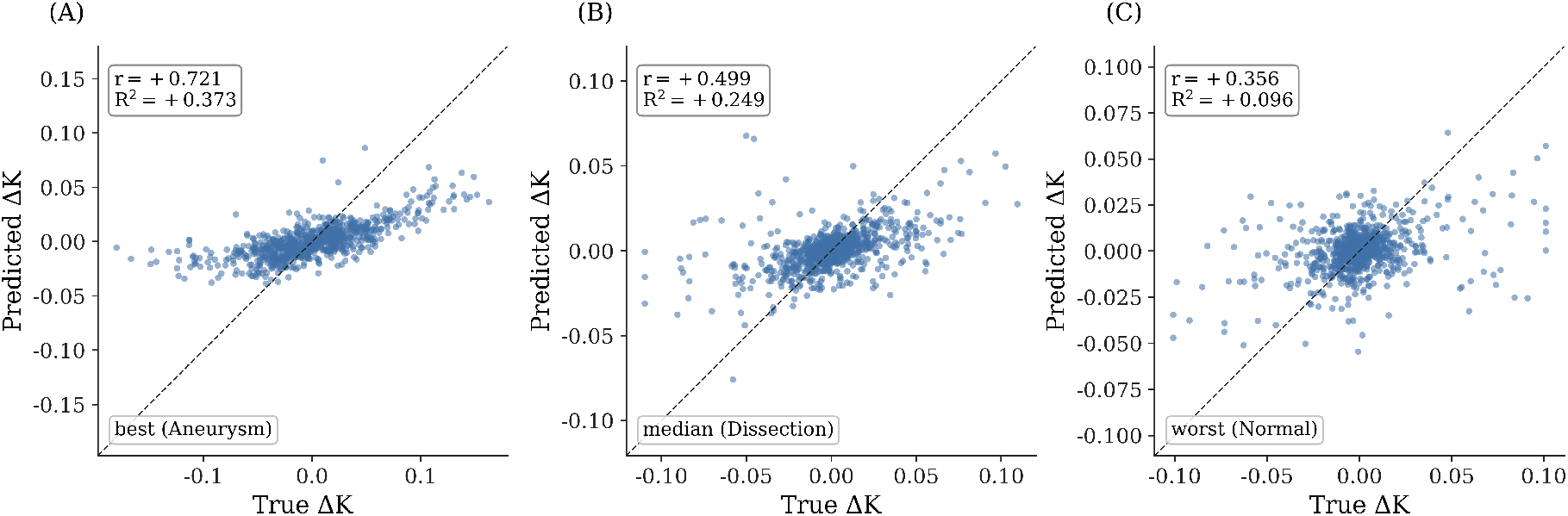
Predicted versus true Δ*K* for representative test surfaces. Three of the *n* = 37 held-out test surfaces ranked by per-patient Pearson *r* are displayed: best (Aneurysm, *r* = +0.721, *R*^2^ = +0.373), median (Dissection, *r* = +0.499, *R*^2^ = +0.249), and worst (Normal, *r* = +0.356, *R*^2^ = +0.096), shown as paired heatmaps of true and predicted Δ*K*. The sign and spatial localization of the largest positive-and negative-Δ*K* patches are visibly recovered on the best and median surfaces; the worst surface preserves spatial localization but compresses the prediction magnitude toward zero, a failure mode consistent with unresolved sub-patch-scale shear.

**Figure 8:**
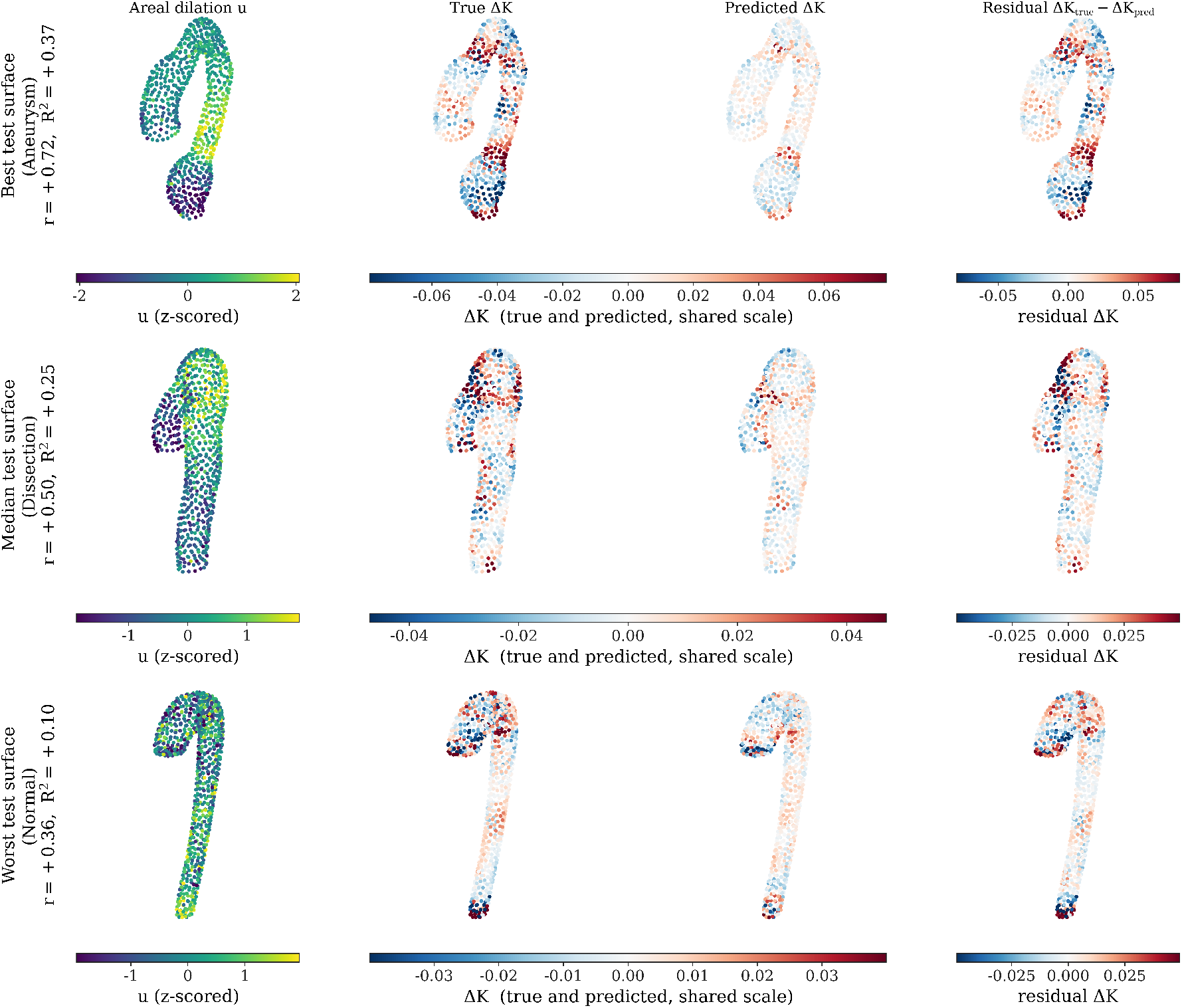
Spatial structure of areal dilation, integrated curvature change, and Multiscale prediction across three test surfaces. Each row corresponds to one held-out test surface, ranked by per-surface Pearson correlation between predicted and true Δ*K*: the best (Aneurysm, *r* = +0.72, *R*^2^ = +0.37), median (Dissection, *r* = +0.50, *R*^2^ = +0.25), and worst (Normal, *r* = +0.36, *R*^2^ = +0.10) test surface. Columns from left to right show the local areal dilation *u* (viridis), the true integrated curvature change Δ*K*, the Multiscale prediction 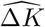, and the residual 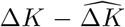. The middle two columns share a symmetric diverging color scale per row so that blue denotes negative and red denotes positive curvature change. Each panel renders the field as a 3D point cloud of patch centroids on the *initial* surface geometry (the t=0 scan, before remodeling), viewed from a consistent camera angle (elev = 20°, azim = 30°); all 800 patches per surface are shown. The coincidence of red and blue regions between true and predicted columns on the best and median rows confirms that the regression preserves the anatomical localization of curvature-change zones; the worst row shows the prediction’s collapse toward an approximately uniform field.

**Figure 9:**
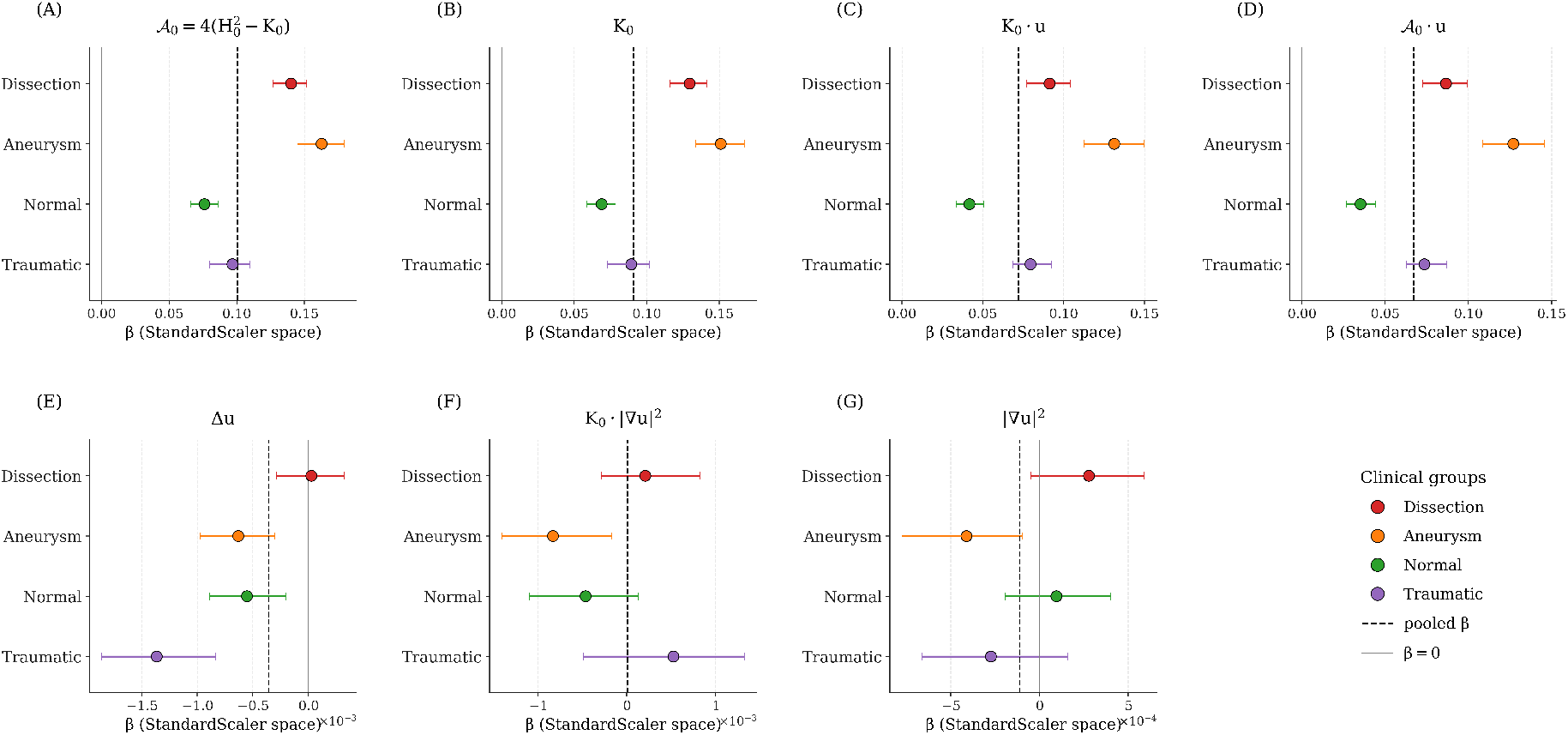
Per-clinical-group sparse-coefficient breakdown for the Multiscale model. The seven sparse-top coefficients are refit separately within each of the four clinical groups (Dissection *n* = 87, Aneurysm *n* = 46, Normal *n* = 41, Traumatic *n* = 25 in the train+val patient subset on which the per-group fit is performed); horizontal errorbars are bootstrap 95% confidence intervals (*B* = 200). Dashed vertical lines mark the pooled-fit *β*. Panels A–D: the four leading local physics terms (𝒜_0_, *K*_0_, *K*_0_*u*,𝒜_0_*u*) are uniformly positive across all four groups with non-overlapping CIs above zero, showing that the headline equation form is shared across physiologic, dissection, aneurysmal, and traumatic geometries. The two anisotropy-driven terms separate the diseased and non-diseased cohorts cleanly: the Aneurysm and Dissection groups carry *β*_*𝒜* 0_ = +0.163 [+0.145, +0.179] and +0.140 [+0.127, +0.152] respectively, while the Normal and Traumatic groups sit at +0.076 [+0.066, +0.086] and +0.097 [+0.080, +0.109], roughly half the diseased magnitude; the same separation is sharper on 𝒜_0_ ·*u*, where the Aneurysm coefficient +0.127 is ∼3.6× the Normal coefficient +0.035. The non-conformal anisotropy correction is therefore quantitatively larger in pathologic remodeling than in physiologic or traumatic geometries, consistent with the shear susceptibility interpretation of 𝒜_0_. Magnitudes follow an Aneurysm ≳ Dissection ≫Traumatic ≳ Normal ordering preserved across all four leading panels: under per-patient demeaning, this is the expected attenuation when within-surface Δ*K* variance is small, so the equation form is shared while the equation magnitude measures how strongly disease-driven remodeling is geometry-organized in each cohort. Panels E–G: the three sub-leading terms (Δ*u, K*_0_ |∇*u*| ^2^, |∇*u*| ^2^) display mixed signs across groups with bootstrap CIs that include zero in most cells, matching the bootstrap-non-significance verdict on the pooled fit and confirming that those terms are correctly identified as below-noise on every group.

Per-patient heterogeneity in *r* tracks per-surface noise scale, since surfaces with intrinsically smaller |Δ*K*| are harder to fit. This motivates the per-patient framing of Section 2.7, and no clinical group shows a systematic floor or ceiling. On a balanced per-patient tercile labeling of Δ*K*, the same predictor returns Cohen’s *κ* = +0.241 on the held-out cohort, with macro *F*_1_ = 0.491 and accuracy = 0.494. The full confusion matrix is in Suppl. Section 13.

## 4 Discussion

The closed-form equation predicts the per-patch change in integrated Gaussian curvature on aortic surfaces directly from the area dilation between two CT scans. On 236 paired thoracic aortas the fit reaches within bootstrap noise of a high-capacity graph neural network ceiling, identifying the unexplained variance as structural shear that two-timepoint registration cannot observe rather than a modeling failure. The synthetic aorta validation supports the equation form on controlled inputs. Under prescribed conformal expansion the fit recovers the analytic Liouville coefficient at every prescribed deformation extent. A focal apex bulge motivates the anisotropy correction, and the proxy 𝒜_0_ = (*κ*_1_ − *κ*_2_)^2^ captures the non-conformal residual that the Liouville triple alone misses. Across a continuous mixture of conformal and non-conformal modes, the anisotropy model holds *R*^2^ ≥ +0.71, with 𝒜_0_ acting as the leading non-conformal correction motivated by morphoelastic shell theory [20–22].

On the cohort, the Liouville source coefficient is positive in nearly every per-patient fit and remains positive in each clinical-group refit. The synthetic conformal value 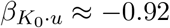 flips to positive on the same idealized arch under a focal non-conformal bulge (Suppl. Section 4, Mode B), and the cohort 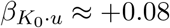 is positive and an order of magnitude attenuated. The anisotropy term enters the cohort fit at magnitudes comparable to the Liouville source.

The anisotropy coefficient *β*_*𝒜* 0_ is roughly twice as large on the diseased cohorts (Aneurysm, Dissection) as on the non-diseased (Normal, Traumatic). The separation is sharper on the coupled term, where the Aneurysm coefficient is about 3.6× the Normal coefficient. Because 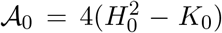 depends only on the initial-scan geometry, the per-patch anisotropy map functions as a shear susceptibility marker computable from a single CT.

The four-level hierarchy ranges from the three-feature Liouville baseline to a graph neural network used as a high-capacity reference. The closed-form Multiscale equation reaches within bootstrap noise of the neural network ceiling. A patch-level variance partition shows that Multiscale recovers about a quarter of the demeaned Δ*K* variance, with the remaining three-quarters structurally unrecoverable from *u, K*_0_, and *H*_0_. Higher-capacity neural network variants overfit the training set yet collapse on held-out test (Suppl. Section 5), identifying the residual as structural rather than a fitting artifact.

The closed-form predictor sits alongside two existing approaches with different trade-offs. Patient-specific morphoelastic finite element simulation reconstructs the full metric evolution, including the unobserved shear, by solving forward from calibrated material parameters and boundary conditions [1, 12]. Deep neural network surrogates trained on vascular geometry, such as LaB-GATr [15], can absorb hemodynamic and prior-growth signals at the cost of a parameter set whose individual entries lack a direct geometric reading. The closed-form equation here uses only imaging-derived quantities, with each coefficient anchored to a specific geometric term in the predictor.

The Liouville bridge could extend to other thin biological surfaces whose evolution carries a substantial conformal component. Cell sheets and bilayers [21, 36] are the closest analog, since morphoelastic shell theory yields the same anisotropy correction as a leading non-conformal term. Cortical surfaces [38, 39] carry an established Ricci-flow tradition that fits the same conformal framework.

The same closed-form structure positions the equation for prospective use beyond the retrospective two-scan setting. With a population model of how the area dilation field evolves between consecutive scans, it could be iterated forward to predict curvature change beyond the second scan; with a population model of the area dilation field conditioned on the initial geometry, it could in principle predict curvature-change distributions from a single baseline scan, without a follow-up. Both extensions require a separate characterization of the population distribution of area dilation, which the present work does not provide.

Two limitations frame future extensions. The local dilation *u* is estimated from two scans [1], with typical cohort |*u*| *<* 0.2 satisfying the small-*u* bound but rapid pathologic dilation potentially violating it. A time-resolved cohort with three or more timepoints would extend the regime and enable a Ricci-flow-coupled estimator using the Liouville equation as a hard constraint [40, 41]. The sub-patch fidelity of non-rigid registration is a known constraint [1, 8], and the synthetic-to-cohort Δ*R*^2^ gap is consistent with sub-patch shear dominating the unexplained variance.

## 5 Conclusion

A closed-form equation derived from the linearized Liouville expansion of the conformal change-of-metric law predicts the change in integrated Gaussian curvature on a biological surface from the imaging-derived area dilation, complementing patient-specific morphoelastic simulation. The predictor adds, to the linearized conformal source, a non-conformal anisotropy proxy that measures the squared difference between principal curvatures of the initial surface and marks where the geometry is susceptible to shear modes predicted by morphoelastic shell theory. On idealized aortic geometries under prescribed conformal expansion, the fitted equation reproduces the analytic Liouville coefficient at every prescribed deformation extent, providing a controlled test of the equation form against a known answer. On the clinical cohort, the same fit reaches the predictive ceiling of a flexible graph neural network, identifying the residual as unobserved deviatoric shear and as a direct measure of how far the *in vivo* growth field departs from conformality. Cohorts imaged at three or more timepoints would extend this construction beyond the linear-time approximation and refine the closed-form description of integrated-curvature evolution on aortic surfaces.

## Ethics statement

All imaging data were obtained under University of Chicago Institutional Review Board approval (IRB20-0653); informed consent for retrospective imaging analysis was waived per IRB protocol, consistent with the Declaration of Helsinki principles for retrospective clinical-imaging research.

## Acknowledgements

We acknowledge the support of the National Institutes of Health, USA, NHLBI, R01HL159205 to LP. The Center for Research Informatics is funded by the Biological Sciences Division, USA at the University of Chicago, with additional funding provided by the Institute for Translational Medicine, CTSA, USA grant number ULITR000430 from the National Institutes of Health. The funders had no role in study design, data collection and analysis, decision to publish, or preparation of the manuscript.

## CRediT author contributions

**Kameel Khabaz**: Conceptualization, Data curation, Methodology, Formal analysis, Investigation, Software, Visualization, Validation, Writing – original draft, Review & editing. **Charlie Davis**: Data curation, Writing – review & editing. **Luka Poč ivaš ek**: Conceptualization, Methodology, Investigation, Visualization, Validation, Writing – original draft, Review & editing, Supervision, Resources, Project administration, Funding acquisition.

## Data availability

Imaging data cannot be deposited in a public repository because they contain protected health information governed by HIPAA; access is governed by the IRB-approved data-availability statements of [1, 4]. Synthetic test geometries used in Section 3.1 are reproducible from a deterministic generation script. All code reproducing the analysis pipeline is available with reasonable request.

