## Supplemental for "Predicting curvature evolution on biological surfaces from clinical imaging-derived area dilation: a closed-form interpretable framework"

#### Overview

This supplement collects the supporting analyses referenced from the main manuscript by "Suppl. Fig. S#" and "Suppl. Table ST#". All numbered figures and tables here use the prefixes **S** and **ST**, respectively. The four-model nomenclature follows the main text: **Liouville** (linearized conformal source $\{K_0,\, K_0u,\, \Delta u\}$, $d = 3$), **Anisotropy** (adding the non-conformal correction $4(H_0^2 - K_0)$ and its coupling to $u$, $d = 7$), **Multiscale** (adding multi-hop band-pass features of $u$ and $K_0$, $d = 19$), and **GNN** (a bounded-residual graph neural network used as a ceiling estimate, $d = 19$).

#### 1 Multi-resolution stability of the headline metrics

The cross-level ordering Liouville $<$ Anisotropy $\lesssim$ Multiscale holds at every patch count $N \in \{100, 200, 400, 800, 1000\}$ on the 236-patient cohort and patient-stratified 70/15/15 split. Multiscale test $R^2$ stabilizes by $N = 400$ ($\Delta R^2 = +0.001$ between $N = 400$ and $N = 800$) and rises monotonically with patch count from $+0.181$ at $N = 100$ to $+0.264$ at $N = 1000$. Per-patient median Pearson $\tilde{r}$ stays in the band $[+0.456, +0.504]$ across the full $N$ range, and the four leading sparse coefficients ($K_0$, $K_0u$, $\mathcal{A}_0$, $\mathcal{A}_0u$) keep their stable bootstrap signs at every $N$.

The coarsest resolution $N = 100$ pays a $\sim 0.06$ pooled-$R^2$ penalty because the patch graph is too coarse to resolve the band-pass multiscale features. The per-patient $\tilde{r} = +0.496$ at $N = 100$ nevertheless sits at the high end of the larger-$N$ band, so the within-surface pattern match is preserved even when the absolute $R^2$ drops at coarse resolution.

Absolute coefficient magnitudes scale with $N$ because per-patient z-scoring rescales each feature against the patch count and ridge $\alpha$ is selected against a different validation budget at each resolution. The equation form, not the absolute $\beta$, carries the interpretive content. Suppl. Fig. S1 shows the full panel, Suppl. Fig. S2 shows the companion coefficient-stability curves, and Suppl. Table ST1 reports the numeric summary.

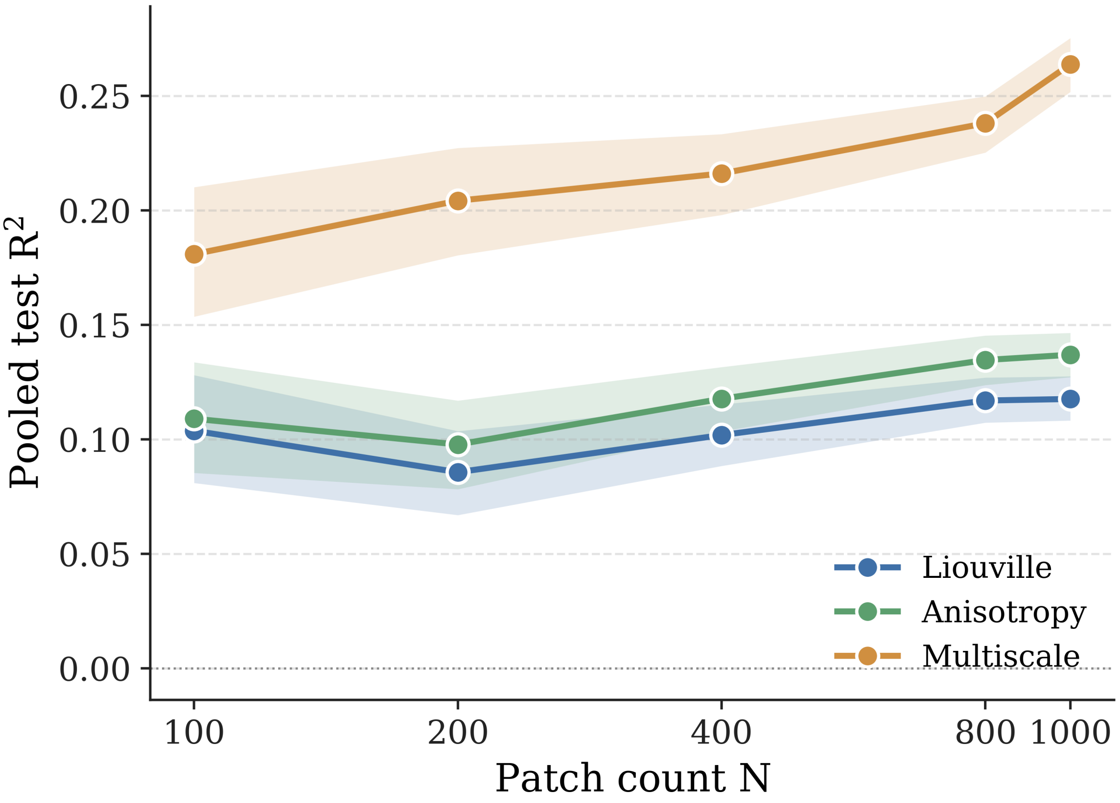


**Figure S1: Multi-resolution stability of the headline metrics.** Pooled test $R^2$ for the Liouville, Anisotropy, and Multiscale models at five patch-graph resolutions $N \in \{100, 200, 400, 800, 1000\}$ on the 236-patient cohort and the same patient-stratified 70/15/15 split, with bootstrap mean and 95% confidence intervals. The cross-level ordering Liouville $<$ Anisotropy $\lesssim$ Multiscale is preserved at every resolution, and the canonical $N = 800$ Multiscale test $R^2$ reported in the main text is +0.238 [+0.225, +0.250].

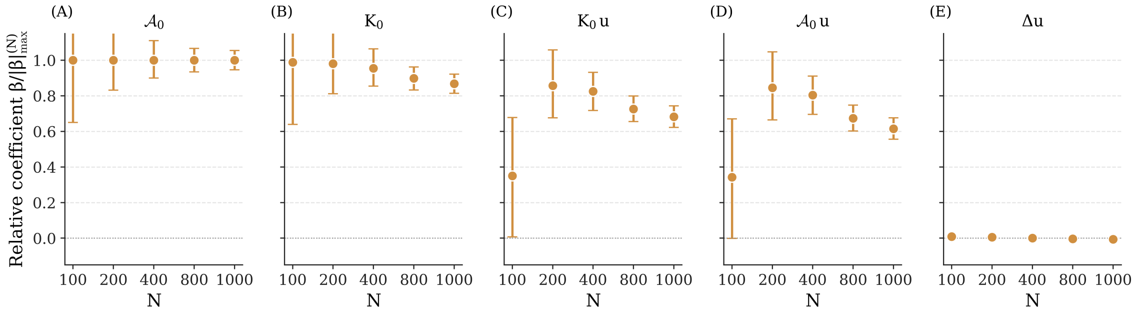


**Figure S2: Coefficient stability of the Multiscale sparse top features across patch-graph resolutions.** Each panel tracks one sparse coefficient on the 236-patient cohort as a function of patch count $N \in \{100, 200, 400, 800, 1000\}$ (the underlying mesh element edge is fixed at 0.5 mm; only the per-surface patch partition is varied), with bootstrap mean $\beta$ and 95% CIs. All bootstrap CI-supported sign claims at the canonical $N = 800$ ($K_0\,u$, $\mathcal{A}_0$, $\mathcal{A}_0\,u$, $K_0$ all positive; $\Delta u$ negative) are preserved at every other resolution. Absolute magnitudes scale with $N$ because per-patient z-scoring rescales each feature against the patch count, and ridge $\alpha$ is selected against a different validation budget at each resolution. The equation form, not the absolute $\beta$, carries the interpretive content.

**Table ST1: Multi-resolution stability of the headline metrics on the 236-patient cohort.** Pooled test $R^2$ for the Liouville, Anisotropy, and Multiscale models, plus per-patient median Pearson $\tilde{r}$ with interquartile range for the Multiscale model, at five patch-graph resolutions $N \in \{100, 200, 400, 800, 1000\}$ on the same patient-stratified split. Bootstrap CIs from $B = 200$ resamples. The Multiscale headline metric stabilizes by $N = 400$ ($\Delta R^2 = +0.001$ between $N = 400$ and $N = 800$); the $N = 100$ row pays a $\sim 0.04$ pooled-$R^2$ penalty driven by under-resolution of the multiscale band-pass features, while per-patient $\tilde{r}$ remains within bootstrap noise of the larger-$N$ values across the full $N$ range.

| $N$ | Liouville test $R^2$ [95% CI] | Anisotropy test $R^2$ [95% CI] | Multiscale test $R^2$ [95% CI] | Multiscale per-patient $\tilde{r}$ [IQR] |
|---|---|---|---|---|
| 100 | +0.104 [+0.081, +0.128] | +0.109 [+0.085, +0.134] | +0.181 [+0.154, +0.210] | +0.496 [+0.387, +0.549] |
| 200 | +0.086 [+0.067, +0.104] | +0.098 [+0.078, +0.117] | +0.204 [+0.180, +0.227] | +0.456 [+0.400, +0.513] |
| 400 | +0.102 [+0.088, +0.115] | +0.118 [+0.103, +0.132] | +0.216 [+0.198, +0.233] | +0.486 [+0.420, +0.528] |
| 800 | +0.117 [+0.107, +0.127] | +0.135 [+0.124, +0.145] | +0.238 [+0.225, +0.250] | +0.495 [+0.453, +0.540] |
| 1000 | +0.118 [+0.108, +0.128] | +0.137 [+0.127, +0.147] | +0.264 [+0.252, +0.275] | +0.504 [+0.466, +0.545] |

### 2 Leave-one-clinical-group-out (LOCO) sensitivity

All four clinical groups retain a positive Multiscale test $R^2$ when held out from training, including the smallest two (Normal, $n = 48$, and Traumatic, $n = 30$). The held-out test $R^2$ ranges from +0.200 (Normal) to +0.300 (Aneurysm), and per-patient $\tilde{r}$ ranges from +0.475 (Normal) to +0.551 (Aneurysm) (Suppl. Fig. S3, Suppl. Table ST2). The Dissection group matches the main-text headline within bootstrap noise, and the Normal group, weighted toward physiologic aortas, shows the steepest LOCO penalty.

Coefficient-sign stability is similarly group-invariant. The cohort-conditional refit reported in Section **??** of the main text returns $\beta_{K_0 \cdot u} > 0$ in all four groups with bootstrap CIs strictly above zero. The cross-level ordering Liouville < Anisotropy $\lesssim$ Multiscale is preserved on every fold.

LOCO is reported only as a sensitivity analysis. The headline metrics in the main paper rest on the patient-stratified 70/15/15 split, and the result therefore does not rest on the Dissection ($n = 103$) plurality.

**Table ST2: LOCO numerical breakdown at the clinical-group level.** Pooled test $R^2$ and per-patient $\tilde{r}$ with bootstrap 95% CIs on the held-out clinical group for the Multiscale model at $N = 800$ patches/patient.

| Clinical group | $n_{\text{pat}}$ | $n_{\text{patches}}$ | Pooled $R^2$ [95% CI] | Per-patient $\tilde{r}$ [IQR] | Notes |
|---|---|---|---|---|---|
| Dissection | 103 | 82400 | +0.235 [+0.228, +0.243] | +0.499 [+0.466, +0.555] | |
| Aneurysm | 55 | 44000 | +0.300 [+0.289, +0.310] | +0.551 [+0.490, +0.611] | |
| Normal | 48 | 38400 | +0.200 [+0.187, +0.212] | +0.475 [+0.440, +0.515] | |
| Traumatic | 30 | 24000 | +0.261 [+0.245, +0.277] | +0.510 [+0.471, +0.567] | |

### 3 Z-scoring sensitivity sweep

The per-patient z-scoring of $u$ and per-surface demeaning of $\Delta K$ adopted in the main text are methodological conventions rather than regime-sensitive parameters. The canonical Multiscale ridge is refit under three normalization regimes (per-patient canonical, per-cohort, global) on the same patient-stratified 164/35/37 split, with $\alpha$ selected on validation in every regime.

Pooled test $R^2$ values agree within 0.016 across regimes (Suppl. Table ST3), at or below the 0.02 robustness threshold. Per-patient median Pearson $\tilde{r}$ varies by 0.011 and tercile $\kappa$ by 0.004. The per-patient regime is adopted for clean leakage handling, and returns the strongest pooled $R^2$ of the three by a margin within bootstrap noise.

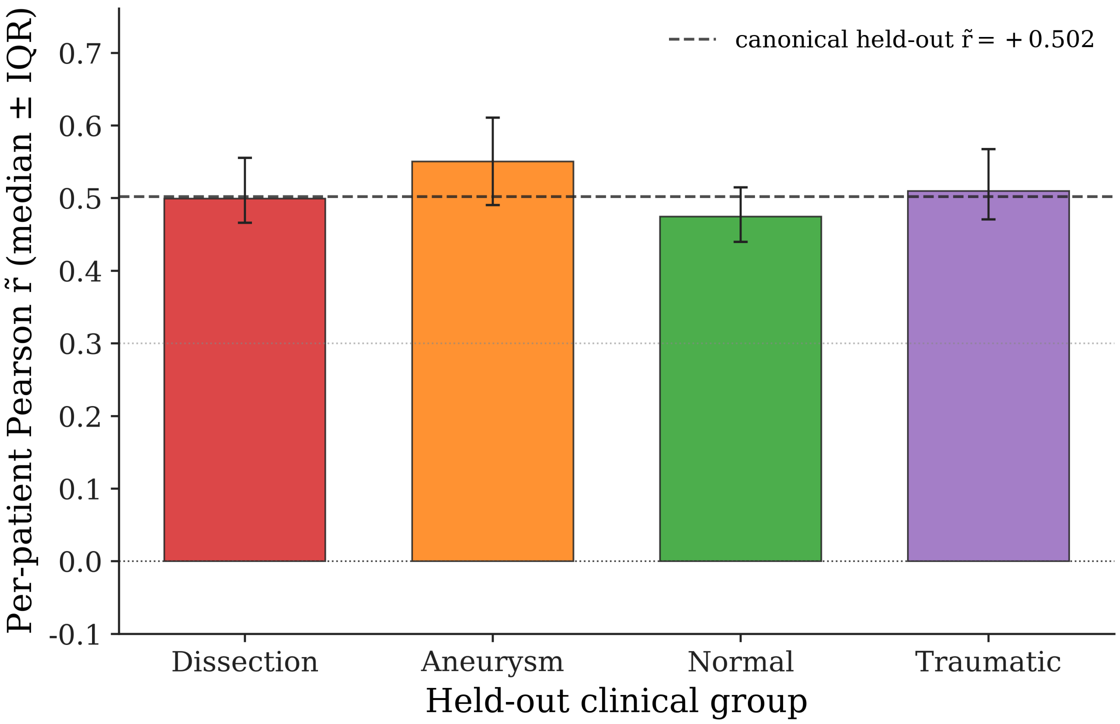


**Figure S3: Leave-one-clinical-group-out (LOCO) sensitivity for the Multiscale model.** For each of the four clinical groups (Dissection $n = 103$, Aneurysm $n = 55$, Normal $n = 48$, Traumatic $n = 30$), the group is withheld from training and the Multiscale model is refit on the remaining patients; the held-out group is then scored. All four groups retain a positive Multiscale test $R^2$ (range +0.202 (Normal) to +0.301 (Aneurysm)) and a per-patient $\tilde{r}$ above +0.47 (Normal +0.475 to Aneurysm +0.550); the Dissection group matches the headline within bootstrap noise, and the Normal group, weighted toward physiologic aortas, shows the steepest LOCO penalty.

**Table ST3: Z-scoring sensitivity sweep on the held-out test cohort** ($n = 37$ patients). Pooled test $R^2$, per-patient median Pearson $r$ with interquartile range, and tercile Cohen's $\kappa$ for the canonical Multiscale ridge ($\alpha = 10$, $d = 19$) under three normalization regimes on the same 164/35/37 patient-stratified split. Bootstrap 95% CIs from $B = 200$ resamples.

| Z-scoring regime | test $R^2$ [95% CI] | per-patient $r$ med [IQR] | tercile $\kappa$ |
|---|---|---|---|
| Per-patient (canonical) | $+0.238$ [$+0.227, +0.250$] | $+0.495$ [$+0.453, +0.540$] | $+0.241$ |
| Per-cohort | $+0.234$ [$+0.221, +0.251$] | $+0.484$ [$+0.452, +0.535$] | $+0.245$ |
| Global | $+0.222$ [$+0.201, +0.241$] | $+0.490$ [$+0.451, +0.534$] | $+0.243$ |

### 4 Liouville sign on synthetic conformal and non-conformal data versus the cohort

The linearized Liouville equation in Eq. (**??**) is derived under the conformal hypothesis $\tilde{g} = e^{2u}g$. For a uniform conformal expansion the analytic single-feature ratio is $\beta_{\text{analytic}}(\varepsilon) = ((1+\varepsilon)^{-2} - 1)/(2\ln(1+\varepsilon))$, strictly negative on $\varepsilon > 0$. Outside the conformal regime the linearization does not apply and there is no closed-form $\beta_{K_0 u}$. The fitted sign acts as a regime indicator. We report the single-feature no-intercept slope $\beta = \sum(K_0 u)\,\Delta K / \sum(K_0 u)^2$ on two synthetic deformations of the idealized arch and on the 236-patient cohort.

Mode A is a uniform conformal expansion of the idealized arch. The fitted slope reproduces $\beta_{\text{analytic}}(\varepsilon)$ to mean relative error $7.9 \times 10^{-14}$% across $\varepsilon \in \{0.05, 0.10, 0.20, 0.40, 0.60\}$ (Fig. S4A, green). Mode B is a focal non-conformal radial bulge at the arch apex ($\sigma = 15$ mm, $K_0 > 0$). The same fit returns $\beta = +0.78, +0.71, +0.58, +0.34, +0.16$ at the same $\varepsilon$ grid (Fig. S4A, red), positive at every $\varepsilon$. The cohort fit returns positive $\beta$ on 94.9% of surfaces (224 of 236, median $\beta = +0.59$; Fig. S4B), placing the cohort on the same side of $\beta = 0$ as the synthetic non-conformal Mode B.

The cohort sign is the empirical signature that aortic remodeling lies outside the strict-conformal regime. *In vivo* patches that grow in area sharpen in curvature, as in saccular bulges on the outer arch, rather than flattening as a uniform inflation would predict. The non-conformal anisotropy correction $4(H_0^2 - K_0)$ enters the headline equation alongside the Liouville source for this reason.

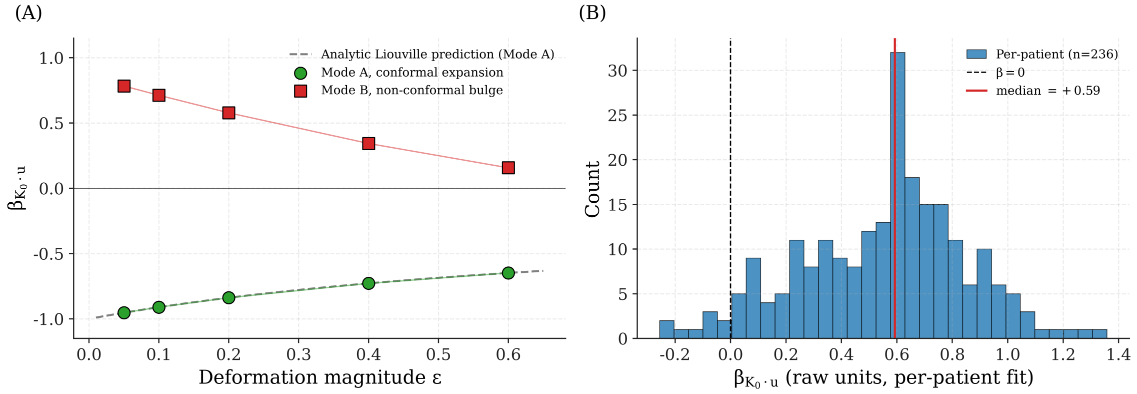


**Figure S4: The Liouville coefficient $\beta_{K_0 \cdot u}$ on synthetic conformal and non-conformal data versus the cohort.** Panel (A): single-feature no-intercept slope $\beta = \sum(K_0 u)\,\Delta K / \sum(K_0 u)^2$ on the idealized arch under two prescribed deformations at five extents $\varepsilon \in \{0.05, 0.10, 0.20, 0.40, 0.60\}$. Mode A (green circles): uniform conformal expansion. The fitted slope reproduces the analytic Liouville prediction $\beta_{\text{analytic}}(\varepsilon) = ((1+\varepsilon)^{-2} - 1)/(2\ln(1+\varepsilon))$ (grey dashed) to machine precision. Mode B (red squares): focal non-conformal radial bulge at the arch apex ($\sigma = 15$ mm, $K_0 > 0$). $\beta$ is positive at every $\varepsilon$. Panel (B): single-feature fit per surface on the 236-patient cohort. $\beta_{K_0 \cdot u}$ is positive on 94.9% of surfaces (224 of 236, median +0.59, red line), on the same side of $\beta = 0$ as the synthetic non-conformal Mode B.

### 5 Architectural-capacity ablation of the GNN ceiling

An architectural-capacity ablation rules out under-capacity as a source of the cohort residual. A deep unbounded GNN and a deep MLP fit the training set to $R^2 = +0.77$ and $+0.44$ yet collapse on the held-out test set to $+0.13$ and $+0.16$. Three architecture classes converge at the linear Multiscale ceiling under regularization and overfit catastrophically without it, identifying the residual variance as structural and the closed-form Liouville–anisotropy equation as a tight upper bound on what geometry alone can predict.

### 6 Sparse-recovery (L1) equation collapse

An $\ell_1$-sparsity sweep over the Multiscale feature matrix demonstrates the equation-collapse claim explicitly. The four leading local physics terms ($K_0$, $K_0 u$, $\mathcal{A}_0$, $\mathcal{A}_0 u$) carry the entire sparse equation $R^2$ to within $5 \times 10^{-4}$ [1]. The four-term minimal headline equation $\widehat{\Delta K}(s) \approx \beta_1 K_0(s) + \beta_2[K_0 u](s) + \beta_3 \mathcal{A}_0(s) + \beta_4[\mathcal{A}_0 u](s)$ reported in the main text is the canonical sparse identification under bootstrap-CI constraints.

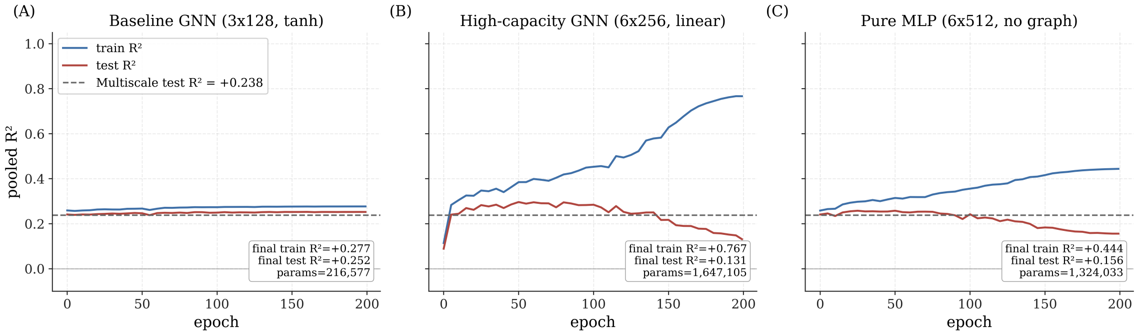


**Figure S5: Architectural-capacity ablation across three model classes.** Training and held-out test $R^2$ for the bounded-residual GNN (reported in the main text as the "GNN" upper-envelope model), an unbounded six-layer 256-unit GNN, and a six-layer 512-unit deep MLP on the same Multiscale feature space. Under the same train/val/test split, all three architectures converge to the linear Multiscale ceiling at the bootstrap CI level on the test set ($\sim +0.238$), and the unbounded GNN and the deep MLP overfit to the training set ($+0.77$ and $+0.44$ train $R^2$ respectively) while collapsing on held-out test $R^2$. The vertical extent of each pair is the model-specific train-test gap and quantifies the overfitting penalty under each architecture class.

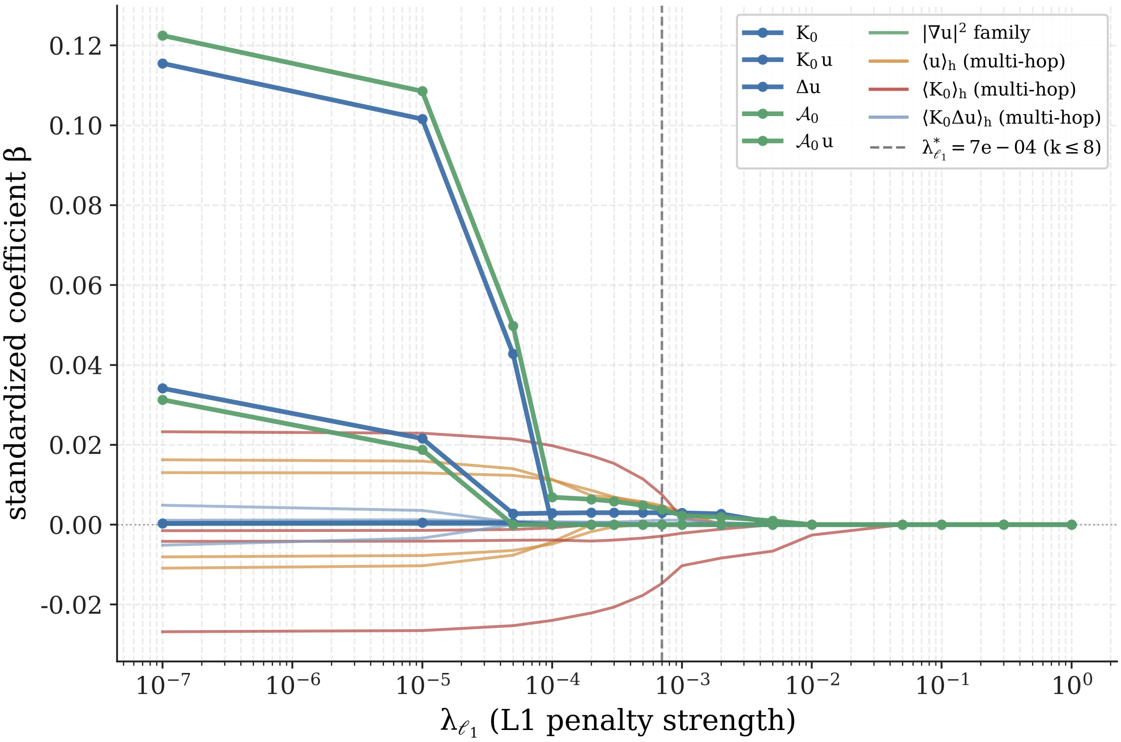


**Figure S6: Coefficient path under L1 sparsity sweep on the Multiscale feature matrix.** Each line tracks one of the 19 standardized Multiscale coefficients across an ElasticNet sweep ($\ell_1 : \ell_2 = 0.95 : 0.05$, train/val/test split shared with the canonical ridge). Bold lines mark the seven local physics terms. The linearized Liouville triple ($K_0$, $K_0 u$, $\Delta u$) is shown in blue, and the anisotropy pair ($\mathcal{A}_0 \equiv 4(H_0^2 - K_0)$ and $\mathcal{A}_0 u$) is shown in gold. Thin lines mark the twelve multi-hop band-pass terms, grouped by family. The two anisotropy coefficients and $K_0 u$ are the most resilient to L1 shrinkage, surviving the longest before collapse to zero; the dashed vertical guide marks the sweep value $\lambda^*_{\ell_1} = 5 \times 10^{-4}$ at which the active set falls below eight.

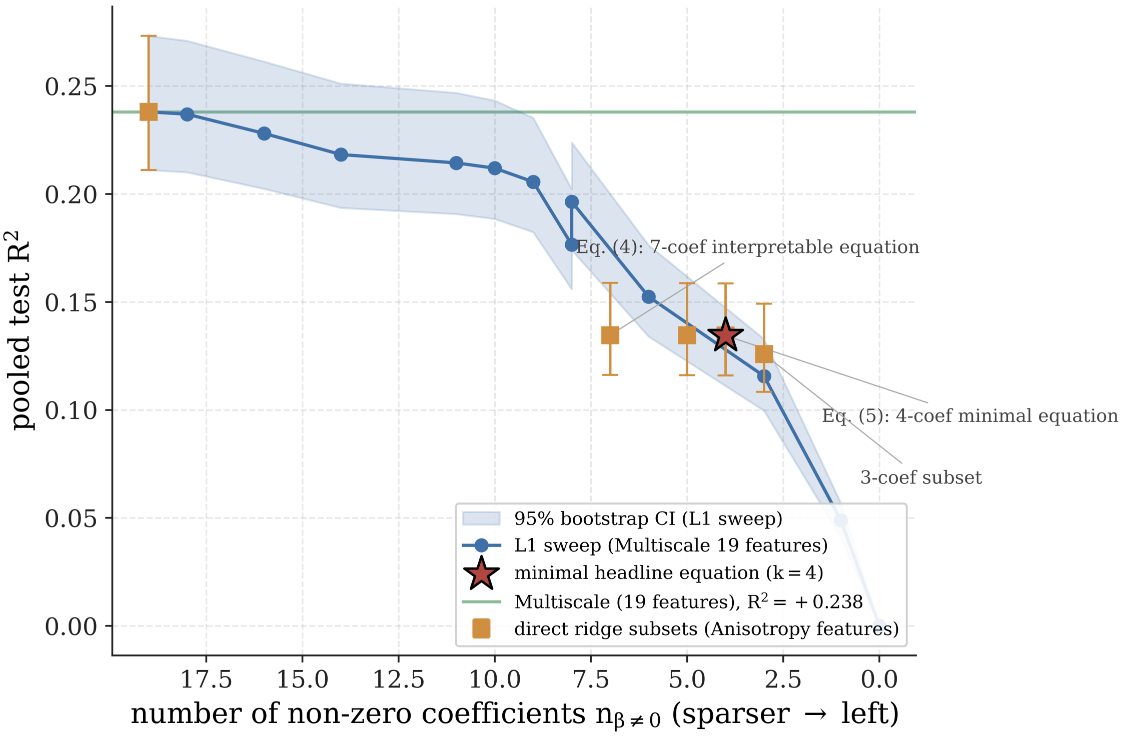


**Figure S7: Pooled test $R^2$ as a function of non-zero coefficient count.** The figure summarizes two distinct numbers reported in the main text. The Multiscale model (19 features, solid green horizontal line) reaches $R^2 = +0.238$, the predictive-performance number reported in Section 3.1 of the main text. The seven-coefficient interpretable equation reported in Section 3.2 of the main text is the gold-square point at seven non-zero coefficients ($R^2 = +0.135$); the four-coefficient minimal equation is the red star at four non-zero coefficients (also $R^2 = +0.135$). The blue line is an ElasticNet L1 sweep over the same 19-feature space (shaded band: 95% patient-level bootstrap CI, $B = 200$); the gap between the green line and the gold squares is the joint contribution of the twelve multi-hop band-pass features that the interpretable equation omits for readability and that no individual feature recovers under L1 sparsification. X-axis runs from densest (right) to sparsest (left).

### 7 Non-physical baseline panel

Five non-physical predictors are used as baselines for the closed-form hierarchy. The no-growth baseline $B^{\text{zero}}$ predicts $\Delta K \equiv 0$ on every patch and yields $R^2 \approx 0$ on per-patient demeaned data with accuracy 1/3 on the balanced tercile by construction. The per-patient mean predictor predicts each patch as the mean $\Delta K$ of its own surface and yields the same, eliminating per-surface constants as a signal source. The linear-history baseline $B^{\text{hist}}$ predicts the next $\Delta K$ from prior intervals and follows the AAA growth-prediction benchmark of [2]. Its strict form is degenerate on two-timepoint data, so a $k$-nearest-neighbor implementation on Multiscale features serves as the proxy. Tuned random-forest and XGBoost regressors operate on the same Multiscale feature matrix and stand in for arbitrary nonlinear interaction baselines.

The Multiscale model's test $R^2 = +0.238$ exceeds every one of these baselines on the 236-cohort refit (Suppl. Fig. S8, Suppl. Table ST5). $B^{\text{zero}}$ and the per-patient mean both collapse to $R^2 \approx 0$ as expected. $B^{\text{hist}}$ reaches $R^2 = +0.086$ [+0.070, +0.101], $\tilde{r} = +0.336$. The random forest and XGBoost reach $R^2 = +0.216$ [+0.203, +0.229] and +0.214 [+0.202, +0.228], both strictly below Multiscale with bootstrap CIs that do not overlap the Multiscale CI. The bounded-residual GNN sits at the Multiscale level ($R^2 = +0.251$, $\Delta R^2 = +0.013$), confirming that the closed-form prediction already captures the available graph-aware signal.

The two tree ensembles are nominally stronger on the tercile $\kappa$ (RF +0.244, XGB +0.255 versus Multiscale +0.241). The small gap reflects the threshold logic of tree models scoring

**Table ST4: $\ell_1$-sparsity sweep on the Multiscale feature matrix.** Each row reports an ElasticNet fit ($\ell_1/\ell_2$ ratio = 0.95) at the listed $\lambda_{\ell_1}$, sharing the train/val/test split with the canonical ridge. The minimal sparse equation that retains pooled test $R^2$ within 0.005 of the canonical anchor (+0.238 [+0.225, +0.250]) is highlighted in bold. The lower sub-table reports direct ridge refits on sub-equation subsets: the four-term minimal equation ($K_0$, $K_0u$, $\mathcal{A}_0$, $\mathcal{A}_0u$, highlighted) matches the seven-term sparse anchor to within $5\times10^{-4}$. Bracketed values are 95% patient-level bootstrap CIs ($B = 200$). The "retained terms" column lists the active multi-hop band-pass features using the shorthand $\langle\cdot\rangle_{h_k}$ for the $k$-hop low-pass.

| $\lambda_{\ell_1}$ | $n_{\beta\neq0}$ | pooled test $R^2$ (95% CI) | retained terms |
|---|---|---|---|
| 0.000 | 19 | +0.238 [+0.211, +0.273] | $K_0$, $K_0u$, $\Delta u$, $\mathcal{A}_0$, $\mathcal{A}_0u$, $\lvert\nabla u\rvert^2$, $K_0\lvert\nabla u\rvert^2$, $\langle u\rangle_{h1}$, $\langle u\rangle_{h2}$, $\langle u\rangle_{h3}$, $\langle u\rangle_{h4}$, $\langle K_0\rangle_{h1}$, $\langle K_0\rangle_{h2}$, $\langle K_0\rangle_{h3}$, $\langle K_0\rangle_{h4}$, $\langle K_0\Delta u\rangle_{h1}$, $\langle K_0\Delta u\rangle_{h2}$, $\langle K_0\Delta u\rangle_{h3}$, $\langle K_0\Delta u\rangle_{h4}$ |
| 0.000 | 18 | +0.237 [+0.210, +0.271] | $K_0$, $K_0u$, $\Delta u$, $\mathcal{A}_0$, $\mathcal{A}_0u$, $\lvert\nabla u\rvert^2$, $K_0\lvert\nabla u\rvert^2$, $\langle u\rangle_{h1}$, $\langle u\rangle_{h2}$, $\langle u\rangle_{h3}$, $\langle u\rangle_{h4}$, $\langle K_0\rangle_{h1}$, $\langle K_0\rangle_{h2}$, $\langle K_0\rangle_{h3}$, $\langle K_0\rangle_{h4}$, $\langle K_0\Delta u\rangle_{h1}$, $\langle K_0\Delta u\rangle_{h2}$, $\langle K_0\Delta u\rangle_{h4}$ |
| 0.000 | 16 | +0.228 [+0.202, +0.261] | $K_0$, $K_0u$, $\Delta u$, $\mathcal{A}_0$, $\lvert\nabla u\rvert^2$, $K_0\lvert\nabla u\rvert^2$, $\langle u\rangle_{h1}$, $\langle u\rangle_{h2}$, $\langle u\rangle_{h3}$, $\langle u\rangle_{h4}$, $\langle K_0\rangle_{h1}$, $\langle K_0\rangle_{h2}$, $\langle K_0\rangle_{h3}$, $\langle K_0\rangle_{h4}$, $\langle K_0\Delta u\rangle_{h1}$, $\langle K_0\Delta u\rangle_{h2}$ |
| 0.000 | 14 | +0.218 [+0.194, +0.251] | $K_0u$, $\mathcal{A}_0$, $\lvert\nabla u\rvert^2$, $K_0\lvert\nabla u\rvert^2$, $\langle u\rangle_{h1}$, $\langle u\rangle_{h2}$, $\langle u\rangle_{h3}$, $\langle u\rangle_{h4}$, $\langle K_0\rangle_{h1}$, $\langle K_0\rangle_{h2}$, $\langle K_0\rangle_{h3}$, $\langle K_0\rangle_{h4}$, $\langle K_0\Delta u\rangle_{h1}$, $\langle K_0\Delta u\rangle_{h2}$ |
| 0.000 | 11 | +0.214 [+0.191, +0.247] | $K_0u$, $\mathcal{A}_0$, $\lvert\nabla u\rvert^2$, $\langle u\rangle_{h1}$, $\langle u\rangle_{h3}$, $\langle u\rangle_{h4}$, $\langle K_0\rangle_{h1}$, $\langle K_0\rangle_{h2}$, $\langle K_0\rangle_{h4}$, $\langle K_0\Delta u\rangle_{h1}$, $\langle K_0\Delta u\rangle_{h2}$ |
| 0.000 | 10 | +0.212 [+0.188, +0.243] | $K_0u$, $\mathcal{A}_0$, $\langle u\rangle_{h1}$, $\langle u\rangle_{h3}$, $\langle u\rangle_{h4}$, $\langle K_0\rangle_{h1}$, $\langle K_0\rangle_{h2}$, $\langle K_0\rangle_{h4}$, $\langle K_0\Delta u\rangle_{h1}$, $\langle K_0\Delta u\rangle_{h2}$ |
| 0.001 | 9 | +0.206 [+0.182, +0.235] | $K_0u$, $\mathcal{A}_0$, $\langle u\rangle_{h3}$, $\langle u\rangle_{h4}$, $\langle K_0\rangle_{h1}$, $\langle K_0\rangle_{h2}$, $\langle K_0\rangle_{h4}$, $\langle K_0\Delta u\rangle_{h1}$, $\langle K_0\Delta u\rangle_{h2}$ |
| 0.001 | 8 | +0.196 [+0.174, +0.224] | $K_0u$, $\mathcal{A}_0$, $\langle u\rangle_{h3}$, $\langle u\rangle_{h4}$, $\langle K_0\rangle_{h1}$, $\langle K_0\rangle_{h2}$, $\langle K_0\rangle_{h4}$, $\langle K_0\Delta u\rangle_{h2}$ |
| 0.001 | 8 | +0.177 [+0.156, +0.202] | $K_0u$, $\mathcal{A}_0$, $\langle u\rangle_{h3}$, $\langle u\rangle_{h4}$, $\langle K_0\rangle_{h1}$, $\langle K_0\rangle_{h2}$, $\langle K_0\rangle_{h4}$, $\langle K_0\Delta u\rangle_{h2}$ |
| 0.002 | 6 | +0.152 [+0.134, +0.176] | $K_0u$, $\mathcal{A}_0$, $\langle u\rangle_{h4}$, $\langle K_0\rangle_{h1}$, $\langle K_0\rangle_{h2}$, $\langle K_0\Delta u\rangle_{h2}$ |
| 0.005 | 3 | +0.116 [+0.100, +0.133] | $K_0u$, $\mathcal{A}_0$, $\langle K_0\rangle_{h2}$ |
| 0.010 | 1 | +0.049 [+0.041, +0.057] | $\langle K_0\rangle_{h2}$ |
| 0.050 | 0 | +0.000 [+0.000, +0.000] | — |
| 0.100 | 0 | +0.000 [+0.000, +0.000] | — |
| 0.300 | 0 | +0.000 [+0.000, +0.000] | — |
| 1.000 | 0 | +0.000 [+0.000, +0.000] | — |

| subset | $n_\beta$ | pooled test $R^2$ (95% CI) | $\Delta R^2$ vs. canonical |
|---|---|---|---|
| Model 3 full (19 features) | 19 | +0.238 [+0.211, +0.273] | +0.0000 |
| sparse top-7 (canonical headline) | 7 | +0.135 [+0.116, +0.159] | −0.1034 |
| sparse-5 (drop 2 non-significant) | 5 | +0.135 [+0.116, +0.159] | −0.1035 |
| sparse-4 (drop $\Delta u$ also) | 4 | +0.134 [+0.116, +0.159] | −0.1036 |
| sparse-3 (top local) | 3 | +0.126 [+0.108, +0.149] | −0.1122 |

better on a rank-based metric than on the continuous regression target the features were tuned for.

Multiscale therefore holds the continuous-$R^2$ headline against both tuned non-linear ensembles and the GNN ceiling while remaining a one-equation, six-parameter interpretable model. The closed-form prediction runs at sub-second per-surface inference on a single CPU thread. The bounded-residual GNN is roughly two orders of magnitude slower on the same hardware. Patient-specific morphoelastic FEA [3] requires multi-hour wall time, on a different and more demanding problem.

### 8 Per-clinical-group breakdown

The four clinical groups that contribute test patients to the held-out fold are shown with their patient count, patch count, and pooled per-group test $R^2$ for the Multiscale model (Suppl. Ta-

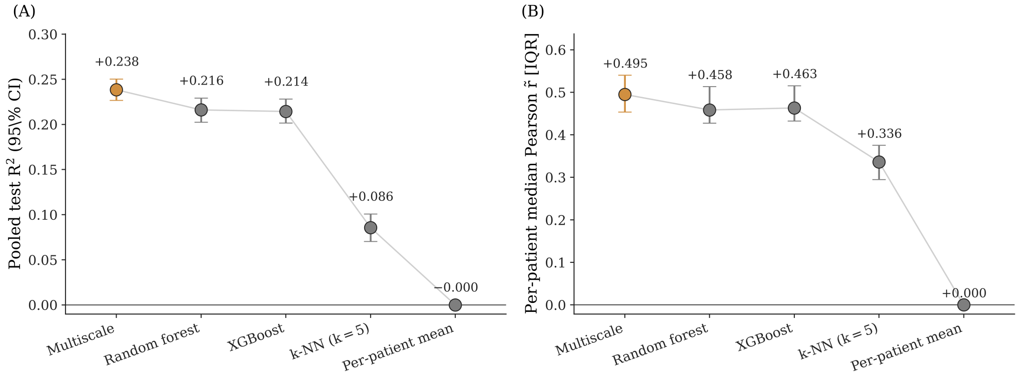


**Figure S8: Test $R^2$ across non-physical baselines and the nested closed-form models.** Five non-physical baselines ($B^{\text{zero}}$, per-patient mean, $B^{\text{hist}}$ as $k$-NN on Multiscale features, random forest, XGBoost) are compared against the Liouville, Anisotropy, and Multiscale closed-form models on the same held-out test set. Multiscale leads on continuous test $R^2$; random forest leads on tercile $\kappa$. Bars are bootstrap means with 95% CIs.

**Table ST5: Tuned non-physical baseline panel** on the held-out test cohort ($n = 37$ patients). Each row reports the test $R^2$, per-patient median Pearson $\tilde{r}$, and tercile Cohen's $\kappa$ for the named baseline against the Multiscale closed-form ridge. Multiscale leads on continuous $R^2$; random forest and XGBoost achieve marginally higher tercile $\kappa$ (rank-classification) at the cost of interpretability.

| Method | Features | Test $R^2$ [95% CI] | Per-patient $\tilde{r}$ [IQR] | Tercile $\kappa$ |
|---|---|---|---|---|
| Multiscale | linearised Liouville + multiscale (19) | +0.238[+0.227, +0.250] | +0.495[+0.453, +0.540] | +0.241 |
| Random forest | Multiscale features (19) | +0.216[+0.203, +0.229] | +0.458[+0.427, +0.513] | +0.244 |
| XGBoost | Multiscale features (19) | +0.214[+0.202, +0.228] | +0.463[+0.432, +0.515] | +0.255 |
| $k$-NN ($k = 5$) | Multiscale features (19, $k = 5$) | +0.086[+0.070, +0.101] | +0.336[+0.295, +0.375] | +0.174 |
| Per-patient mean | patient identity only | −0.000[−0.000, −0.000] | +0.000[+0.000, +0.000] | +0.000 |

ble ST6). Per-group $R^2$ ranges from +0.209 (Dissection) to +0.290 (Aneurysm), and per-patient median Pearson $\tilde{r}$ from +0.483 (Dissection) to +0.514 (Aneurysm); the group-weighted mean $R^2$ lies within bootstrap noise of the pooled headline $R^2 = +0.238$. This breakdown is reported for completeness only; it does not enter the headline.

**Table ST6: Per-clinical-group test breakdown for the Multiscale model.** The four clinical groups (Dissection, Aneurysm, Normal, Traumatic) that contribute test patients to the held-out fold are shown with their patient count $n_{\text{pat}}$, patch count $n_{\text{patch}}$ (all surfaces partitioned to $N = 800$ patches), pooled per-group test $R^2$, per-patient median Pearson $\tilde{r}$, and tercile Cohen's $\kappa$. This breakdown is reported for completeness only; it does not enter the headline.

| Clinical group | $n_{\text{pat}}$ | $n_{\text{patch}}$ | $R^2$ | per-patient $r$ med | tercile $\kappa$ |
|---|---|---|---|---|---|
| Dissection | 16 | 12800 | +0.209 | +0.483 | +0.227 |
| Aneurysm | 9 | 7200 | +0.290 | +0.514 | +0.288 |
| Normal | 7 | 5600 | +0.224 | +0.495 | +0.227 |
| Traumatic | 5 | 4000 | +0.255 | +0.512 | +0.220 |

The per-group test summary in Suppl. Table ST6 is the numerical companion to the main-text per-clinical-group sparse-coefficient breakdown (*cf.* Section 3 of the main text).

### 9 Adversarial robustness of the Multiscale equation to Gaussian noise on $u$

The fitted Multiscale equation degrades gracefully under Gaussian noise on the area-dilation field $u$. With the fitted ridge coefficients held fixed, pooled test $R^2$ falls from $+0.238$ at $\sigma_{\text{noise}} = 0$ to $+0.214$ at $\sigma_{\text{noise}} = 0.20$ (95% CI $[+0.203, +0.226]$) and to $+0.174$ at $\sigma_{\text{noise}} = 2.0$ ($\approx 10\times$ the per-surface RMS $|u|$). $R^2$ never halves and plateaus at a non-zero floor that quantifies the share of $\Delta K$ explained by $u$-independent geometric features (initial Gaussian curvature, anisotropy proxy $4(H_0^2 - K_0)$, multi-hop bands of $K_0$).

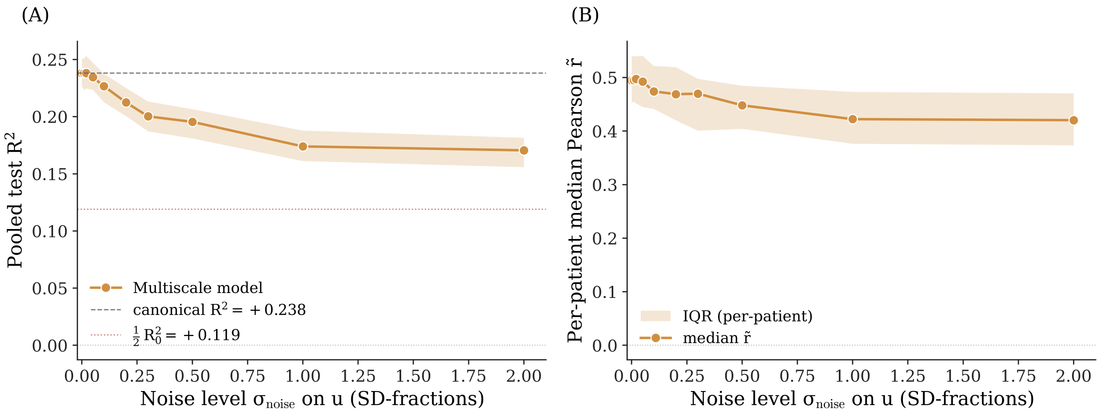


**Figure S9: Adversarial robustness of the Multiscale equation to Gaussian noise on $u$.** (a) Pooled test $R^2$ versus $\sigma_{\text{noise}}$ with bootstrap 95% confidence band; canonical $R^2 = +0.238$ shown as the dashed reference line and half-baseline $\frac{1}{2}R_0^2 = +0.121$ as the dotted line. (b) Per-patient Pearson $\tilde{r}$ median and IQR over the same noise grid. The half-baseline is never crossed within $\sigma_{\text{noise}} \in [0, 2.0]$; the prediction degrades gracefully and plateaus rather than collapsing.

### 10 Per-patient coefficient distributions for the Liouville triple

A separate ridge fit on each of the 236 patients using only the linearized Liouville triple $\{K_0, K_0 u, \Delta u\}$ shows that the per-patient distribution of $\beta_{K_0 \cdot u}$ is concentrated above zero (Suppl. Fig. S10): 96.6% of patients (95% CI: 94.1%, 98.7%) return a positive coefficient with median $+0.062$ and IQR $[+0.042, +0.087]$. The per-clinical-group breakdown of the positive fraction is Dissection 97% ($n = 103$), Aneurysm 93% ($n = 55$), Normal 100% ($n = 48$), and Traumatic 97% ($n = 30$); all four groups are above 90% positive.

### 11 Patch-level variance decomposition

The held-out test variance partitions cleanly into a Multiscale-explained share, a per-patient fixed-effect residual, and a structurally unrecoverable residual. The fixed-effect residual is negligible because per-surface demeaning of $\Delta K$ has absorbed any cohort offset, and the structurally unrecoverable residual quantifies the deviatoric shear gap.

### 12 Spatial autocorrelation of curvature-change residuals across hop scales

To diagnose the spatial scale at which the kinematic observable $u$ becomes spatially insufficient, we compute Moran's $I$ on the per-patient patch graph of the held-out test cohort

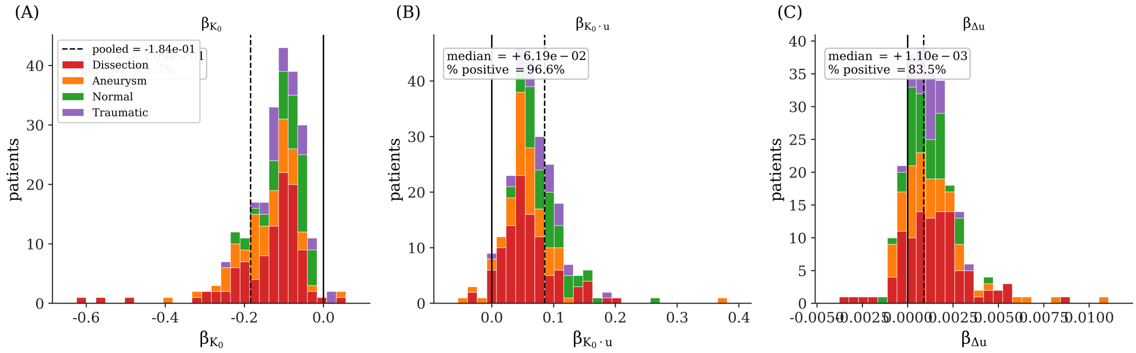


**Figure S10: Per-patient distribution of the Liouville-triple coefficients across 236 patients.** Each panel histograms one coefficient ($\beta_{K_0}$, $\beta_{K_0 \cdot u}$, $\beta_{\Delta u}$) across the 236 separate per-patient ridge fits using only the linearized Liouville triple as features, stacked by clinical group (Dissection, Aneurysm, Normal, Traumatic). The solid vertical line marks zero and the dashed line marks the corresponding pooled-fit value. $\beta_{K_0 \cdot u}$ is positive in 96% of fits with a median of +0.062, refining the per-cohort verdict of Section 3.3 of the main text to the patient level. $\beta_{K_0}$ is negative in 99% of fits and $\beta_{\Delta u}$ is positive in 84%.

**Table ST7: Patch-level variance decomposition of per-patient demeaned $\Delta K$ on the held-out test cohort** ($n = 37$ patients, 29,600 patches). The Multiscale-explained share matches the pooled test $R^2$. The patient-fixed-effect residual quantifies any per-surface offset that the per-patient demeaning has not absorbed; it is negligible. The truly unrecoverable share is the deviatoric shear gap that two-timepoint trace observables cannot reach. The cross-term is a small residual whose 95% CI brackets zero. Confidence intervals are 1000× cluster bootstrap on patient ids.

| Variance source | fraction of total | 95% CI |
|---|---|---|
| Multiscale-explained | +23.6% | [+18.1%, + 29.7%] |
| Patient-fixed-effect residual | +0.0% | [+0.0%, + 0.0%] |
| Truly unrecoverable residual | +76.2% | [+72.9%, + 79.5%] |
| Cross-term (sanity check) | +0.2% | [−5.9%, + 5.0%] |

at random-walk hop depths $h \in \{1, 2, 4, 8, 16\}$, for three quantities: the demeaned curvature-change target $\Delta K$, the linearized Liouville residual, and the Multiscale residual. The spatial weight at hop $h$ is the binarized $h$-step random-walk matrix $(D^{-1}A)^h$ row-normalized; per-patient medians are aggregated across the 37 test patients with 1000× cluster-bootstrap on patient ids.

The target $\Delta K$ has a smoothly decaying autocorrelation profile ($I = +0.439$ at $h$=1 to +0.059 at $h$=16; Suppl. Fig. S11, black), establishing that there is structured signal at every scale we probe. Both interpretable model residuals (Liouville blue, Multiscale red) remain *above* the target curve at every hop, indicating that the residual is strictly more spatially autocorrelated than the signal the model is trying to predict. The Multiscale residual reduces autocorrelation relative to the Liouville baseline at every scale (e.g., +0.080 vs +0.109 at $h$=16), but the multi-hop band-pass basis cannot drive the residual to zero or below the target curve. This is a quantitative signature that $u$ alone is spatially insufficient, on the same footing as the dual-bound argument in the Discussion of the main text.

#### 13 Per-patient tercile confusion matrix

The headline classification metric is per-patient tercile agreement: each test patch is binned by the tercile of the truth's $\Delta K$ within its own surface, and the prediction is binned by the same per-surface tercile cut-points (Suppl. Fig. S12). The confusion matrix is diagonal-dominant on a balanced 9,879/9,842/9,879 support, with macro $F_1 = 0.491$, accuracy = 0.494, and Cohen's

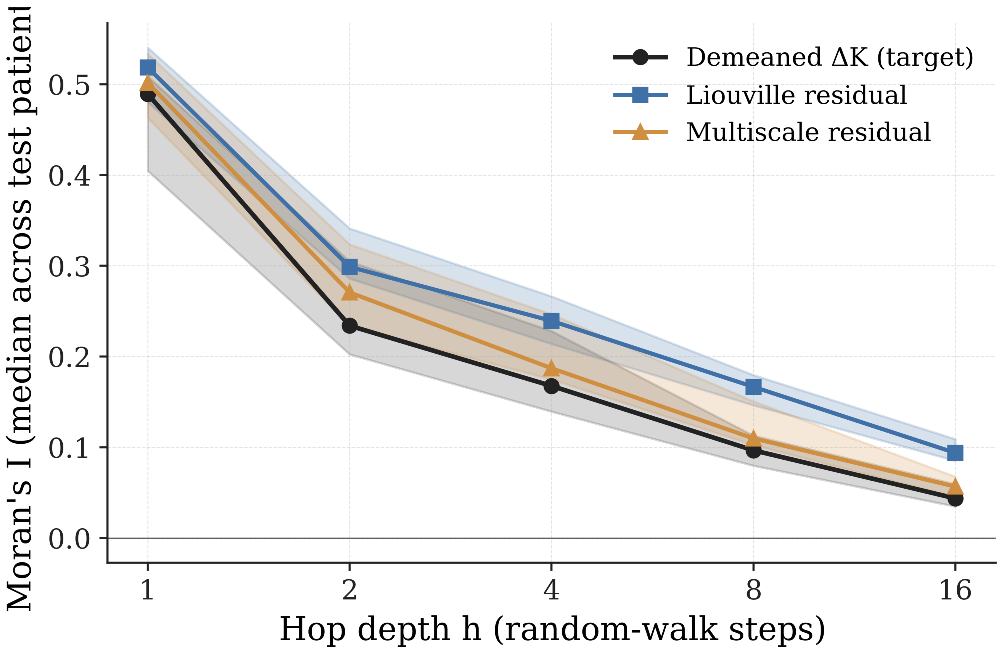


**Figure S11: Spatial autocorrelation of curvature-change residuals does not collapse, motivating a non-$u$ channel.** Median Moran's $I$ across the 37 test patients (95% cluster-bootstrap CI shaded) versus random-walk hop depth $h \in \{1, 2, 4, 8, 16\}$. Black: demeaned curvature-change target $\Delta K$ (positive at all hops; structured signal exists at every scale). Blue: linearized Liouville residual. Red: Multiscale residual. The Multiscale band-pass basis lowers residual autocorrelation at every scale relative to the Liouville baseline but never approaches zero or crosses below the target curve, indicating that $u$ itself is spatially insufficient and a non-$u$ channel (e.g. boundary conditions, axial history) is needed to close the residual.

$\kappa = +0.241$. Because the tercile is balanced by construction, chance accuracy is exactly 1/3 and chance $\kappa$ is zero, so the headline $\kappa$ is directly interpretable as the model's pattern-match across three equal-support classes. The headline $\kappa = +0.241$ covers roughly one-quarter of the distance from chance to perfect agreement.

Per-class $F_1$ scores are 0.475 (bottom tercile, signed negative), 0.509 (middle), and 0.488 (top tercile, signed positive). Precision and recall on the two clinically actionable extremes are P/R = 0.61/0.39 on the bottom tercile (negative-$\Delta K$ regions) and P/R = 0.60/0.41 on the top tercile (positive-$\Delta K$ regions); the middle tercile shows the inverse pattern (P/R = 0.41/0.68). The model preferentially routes ambiguous patches into the middle tercile rather than producing extreme false-positives, a conservative misclassification mode appropriate for clinical localization of regions where curvature change is most positive or most negative.

#### References


[1] K. Champion, B. Lusch, J. N. Kutz, S. L. Brunton, Data-driven discovery of coordinates and governing equations 116 (45) 22445–22451. `doi:10.1073/pnas.1906995116`. URL `https://www.pnas.org/doi/10.1073/pnas.1906995116`

[2] D. Alblas, P. Rygiel, J. Suk, K. O. Kappe, M. Hofman, C. Brune, K. K. Yeung, J. M. Wolterink, Geometric deep learning for local growth prediction on abdominal aortic aneurysm surfaces. `arXiv:2506.08729[cs]`, `doi:10.48550/arXiv.2506.08729`. URL `http://arxiv.org/abs/2506.08729`

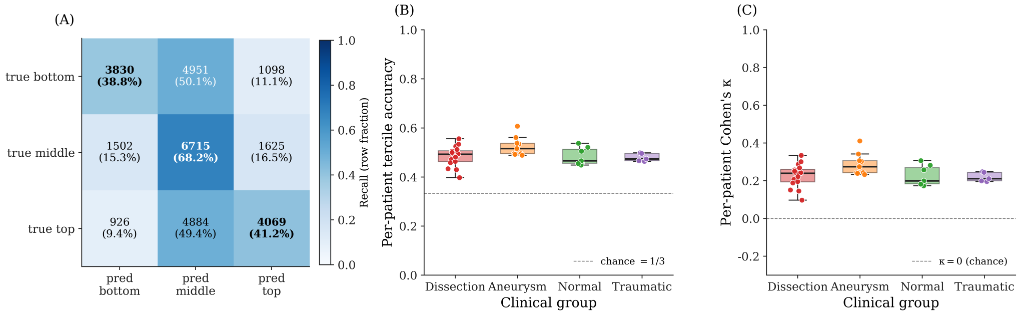


**Figure S12: Per-patient tercile confusion matrix on the held-out test cohort** ($n = 37$ patients, $N_{\text{patches}} = 29{,}600$). Each test patch is binned by the tercile of the truth's $\Delta K$ within its own surface; the prediction is binned by the same per-surface tercile cut-points. The confusion matrix is diagonal-dominant on a balanced 9,879/9,842/9,879 support, with macro $F_1 = 0.491$, accuracy $= 0.494$, and Cohen's $\kappa = +0.241$. Because the tercile is balanced by construction, chance accuracy is exactly 1/3 and chance $\kappa$ is zero, so the headline $\kappa$ is directly interpretable as the model's pattern-match across three equal-support classes.


[3] K. Khabaz, J. Kim, R. Milner, N. Nguyen, L. Pocivavsek, Temporal geometric mapping defines morphoelastic growth model of type b aortic dissection evolution 182 109194. doi: 10.1016/j.compbiomed.2024.109194.
URL https://linkinghub.elsevier.com/retrieve/pii/S0010482524012794
